# Disruption of molecular interactions between G3BP1 stress granule host protein and nucleocapsid (NTD-N) protein impedes SARS-CoV-2 virus replication

**DOI:** 10.1101/2024.10.27.620470

**Authors:** Preeti Dhaka, Ankur Singh, Sanketkumar Nehul, Shweta Choudhary, Prasan Kumar Panda, Gaurav Kumar Sharma, Pravindra Kumar, Shailly Tomar

**Author notes:** corresponding authors, ***Corresponding authors *Shailly Tomar**- Professor, Department of Biosciences and Bioengineering, Indian Institute of Technology Roorkee, Uttarakhand (247667), India;, ***Pravindra Kumar-** Professor, Department of Biosciences and Bioengineering, Indian Institute of Technology Roorkee, Uttarakhand (247667), India;, *Gaurav Kumar Sharma, Senior scientist, Indian Veterinary Research Institute, Izatnagar, Bareilly, Uttar Pradesh state (243122), India. Equal contribution in this manuscript (Preeti Dhaka, Ankur Singh).

## Abstract

The Ras GTPase-activating protein SH3-domain-binding protein 1 (G3BP1) serves as a formidable barrier to viral replication by generating stress granules (SGs) in response to viral infections. Interestingly, viruses, including SARS-CoV-2, have evolved defensive mechanisms to hijack SG proteins like G3BP1 for the dissipation of SGs that lead to the evasion of host’s immune responses. Previous research has demonstrated that the interaction between the NTF2-like domain of G3BP1 (G3BP1_NTF-2_) and the intrinsically disordered N-terminal domain (NTD-N_1-25_) of the N protein plays a crucial role in regulating viral replication and pathogenicity. Interestingly, the current study identified an additional upstream stretch of residues (128KDGIIWVATEG138) (N_128-138_) within the N-terminal domain of the N protein (NTD-N_41-174_) that also forms molecular contacts with the G3BP1 protein, as revealed through *in silico* analysis, site-directed mutagenesis and biochemical analysis. Remarkably, WIN-62577, and fluspirilene, the small molecules targeting the conserved peptide binding pocket in G3BP1_NTF-2,_ not only disrupted the protein-protein interactions (PPIs) between the NTD-N_41-174_ and G3BP1_NTF-2_ but also exhibited significant antiviral efficacy against SARS-CoV-2 replication with EC_50_ values of ∼1.8 µM and ∼1.3 µM, respectively. The findings of this study, validated by biophysical thermodynamics and biochemical investigations, advance the potential of developing therapeutics targeting the SG host protein against SARS-CoV-2, which may also serve as a broad-spectrum antiviral target.

## INTRODUCTION

Viral infection and replication trigger antiviral innate immune responses in the infected host cells. In response, host cells produce cytoplasmic granular aggregates known as stress granules (SGs). However, viruses can hijack these SGs to facilitate their replication, contributing to later-stage disease progression.^1,2^ Viral genome replication is established by the direct interactions of viral proteins with various cellular host proteins, which helps the virus to evade the host’s immune responses and support the viral life cycle.^3,4^ Previous reports showed that several positive single-stranded viruses like Severe acute respiratory syndrome coronavirus 2 (SARS-CoV-2)^5–11^, Chikungunya virus (CHIKV)^12–14^, Zika virus (ZIKV)^15,16^, Dengue virus (DENV)^17–20^, Mouse hepatitis virus (MHV)^5,21,22^ Hepatitis C virus (HCV)^23–25^, Semliki Forest virus (SFV)^26–28^, Sindbis virus (SINV)^29–32^, Encephalomyocarditis Virus (EMCV)^33^, and the Poliovirus (PV)^34–37^ utilize the components of SGs for their own viral replication mechanism to escape from the host immune defence system.^38,39^ The SG assembly formation includes the Ras GTPase-activating protein-binding protein 1 (G3BP1)^40^, G3BP2^41^, T-cell intracellular antigen-1 (TIA-1)^42^, and Caprin1^43^, which protect the host cell genome after virus infection.^44^ The key players of SGs, G3BP1/2 host proteins, directly interact with the N-terminal domain of nucleocapsid protein (NTD-N_41-174_) and support SARS-CoV-2 infection. These host-virus protein-protein interactions (PPIs) destabilize the SG assembly.^45–48^ Interactome investigations suggested that G3BP1/2 are the critical host stress granule intercommunicators for the SARS-CoV-2 infection.^49,50^ The SG assembly proteins generally protect the cell environment against viral infections. However, the NTD-N protein of SARS-CoV-2 rewires the target NTF2-like domain of G3BP1 (G3BP1_NTF-2_) and controls the host cell machinery, destabilizing SG assembly.^11,51–53^ Therefore, G3BP1_NTF-2_, the host cell SGs component, is a promising antiviral target because of its essential role in specific PPIs with the viral proteins utilizing the NTD-N protein.^54,55^ PPIs are potential targets for drug discovery in various diseases such as cancer/degenerative diseases and viral infections, including SARS-CoV-2 coronavirus.^56,57^

Coronaviruses belong to the *coronaviridae* family and the genus Betacoronavirus.^58^ SARS-CoV-2, a single-stranded positive-sense RNA virus, was an accountable reason for the COVID-19 pandemic.^59^ The recurrent bouts of infections due to emerging variants of SARS-CoV-2, being still reported worldwide, are an ominous sign for global health.^60^ The possibility of the reoccurrence of highly transmissible, more virulent, and vaccine-resistant strains in the future continues to be a concern.^61^ Therefore, it is important to develop effective antiviral therapeutics with potential host antiviral targets, as it is very likely that the host-directed antiviral molecule will be broad spectrum and more effective towards the virus mutants.^62–64^

The SARS-CoV-2 proteins manipulate the host cellular processes and enhance its replication and infection.^65–67^ SARS-CoV-2 contains four structural proteins: spike (S), envelope (E), matrix (M), and N protein.^68^ The N protein is amenable to the compaction of viral genomic RNA within the virion, and it induces the formation of the ribonucleoprotein (RNP) complexes via the phase separation process.^69,70^ N protein consists of two functional domains interconnected by intrinsically disordered regions (IDRs). The N-terminal domain (NTD-N_41-174_) and C-terminal domain (CTD-N_247-365_) are responsible for establishing direct contact with the RNA and are involved in the oligomerization process, respectively^71,72^. The intervening IDRs, including IDR-1_1-40_ (N-arm), also interacted with RNA and G3BP1_NTF-2_ protein. At the same time, IDR-2_175-246_ constitute the linker region that connects both NTD with CTD, and the last one, IDR-3_366-409_ forms a C-tail after CTD, as shown in Figure 1(a).^73,74^

**Figure 1.**
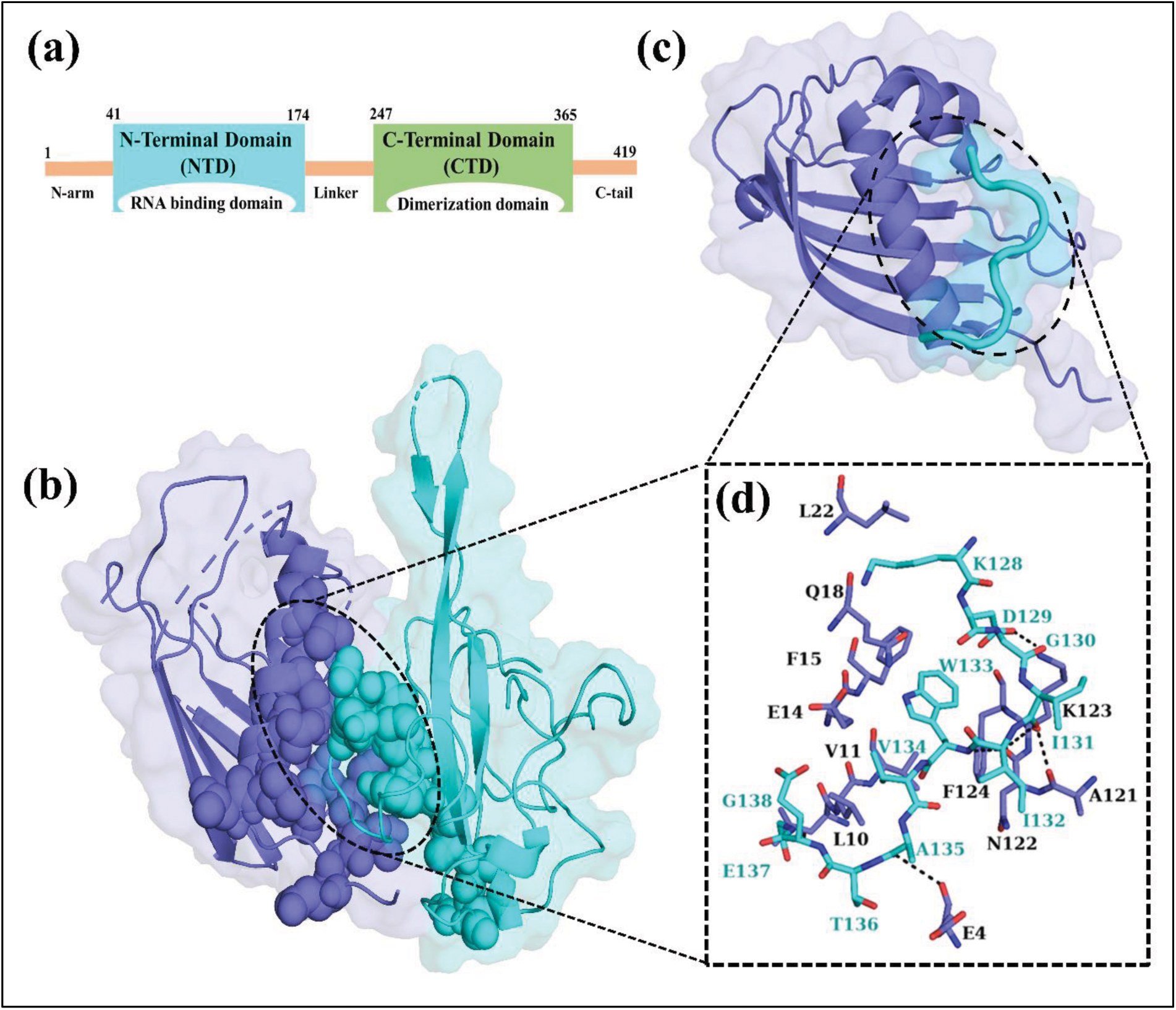
Overview of SARS-CoV-2 N protein and PPIs analysis between NTD-N_41-174_ and the G3BP1_NTF-2_. (a) The N protein consists of two functional domains, NTD (cyan) and CTD (green), separated by an intrinsically disordered region (IDR) linker, flanked by additional IDRs at the N-arm and C-tail (orange). (b) Molecular docking of NTD-N_41-174_ (cyan) with the G3BP1NTF-2 (blue) protein to analyze the PPIs site (Black dashed circle with spheres). (c) Molecular docking of NTD-N_41-174_ derived (128KDGIIWVATEG138) or (N128-138) peptide (cyan cartoon) on G3BP1NTF-2 (blue) encircled with black dashes. (d) Key interacting residues N_128-138_ peptide (cyan sticks) are represented in the insight box, showing interactions with conversed residues of G3BP1NTF-2 (blue sticks).

The IDR-1 and the NTD-N_41-174_ of N-protein are involved in molecular contacts with the NTF2-like domain of the G3BP1_NTF-2_ host protein.^40,48,51^ The detailed molecular interactions with G3BP1_NTF-2_ at the PPIs interface have been reported by determining the crystal structure of the N-terminal peptide (N_1-25_) of the IDR-1 in complex with the G3BP1_NTF2_ (PDB: 7SUO^51^).^70,74^ The peptide N_1-25_ interacts with a conserved pocket in the NTF2-like domain of G3BP1_NTF-2_ that has been extensively studied for its role in binding to other viral proteins and also to the host protein Caprin1.^70^ Likewise, the crystallographic structure of the G3BP1_NTF-2_ with a peptide from the non-structural protein 3 (nsP3) of alphavirus also demonstrates very similar molecular interaction mechanism at the PPI interface.^54,75^ Therefore, the structural comparison of G3BP1_NTF-2_ in complex with nsP3 of alphavirus (PDB: 5FW5)^54^ and the nucleocapsid peptide (NTD-N_1-25_) of SARS-CoV-2 (PDB: 7SUO)^51^ demonstrates that certain viruses commonly utilize the G3BP1_NTF-2_ to establish specific PPIs between the viral proteins and the host protein to surmount the antiviral responses (Figure 2). These PPIs between viral proteins and host G3BP1_NTF-2_ are crucial for viral evasion from host immune responses.^76–78^ The antiviral molecules against CHIKV that target the nsP3 binding pocket in the G3BP1 protein have recently been reported.^55,79^ This means that the interacting interface between the host protein G3BP1_NTF-2_ and viral protein NTD-N_41-174_ is a potential broad-spectrum antiviral target. Identifying potential molecules blocking this PPI and inhibiting SARS-CoV-2 is expected to open new avenues for host-based antiviral drug discovery.

**Figure 2.**
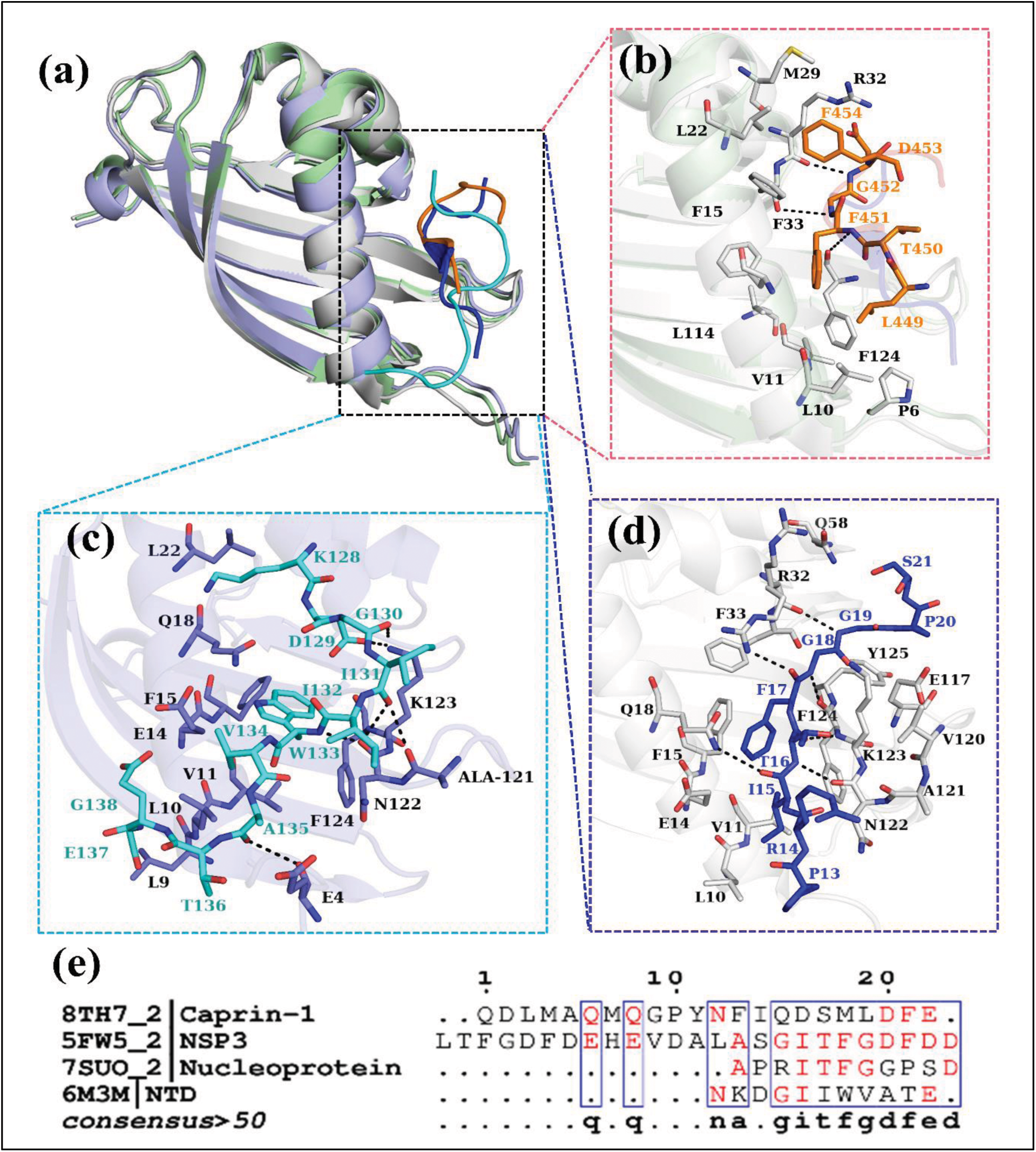
Superimposition of crystal structures of G3BP1-nsP3 (PDB: 5FW5)^54^ and G3BP1 complexed with N_1-25_-IDR region of N-protein (PDB: 7SUO)^51^ and docked structure of the upstream NTD-N_41-174_ derived peptide (N_128-138_) with G3BP1_NTF-2_. (a) The cartoon illustrations of the superimposed PDBs; 5FW5 (green/blue), and 7SUO (grey), represent the conserved site of the G3BP1_NTF-2_ complex structures (on left side). (b) The FGDF motif of nsP3 protein (orange), (c) NTD- N_128-138_ peptide (128KDGIIWVATEG138) (cyan sticks), and (d) the N_1-25_-IDR region of N-protein (12APRITFGGPSD22) (dark blue) are represented in the insight boxes, while the conversed residues of G3BP1_NTF-2_ (light green/white/light blue sticks). Black-coloured dotted lines show the intermolecular H-bond interactions. (e) Sequence alignment of different peptides from host protein Caprin1 (PDB: 8TH7) and different viruses including nsP3 of SFV (PDB: 5FW5), APRITFGGPSD motif the SARS-CoV-2 NTD N_1-25_ N-protein (PDB: 7SUO), and SARS-CoV-2 NTD-N_41-174_ derived peptide (N_128-138_) (PDB: 6M3M), that interacts with conserved pocket of G3BP1_NTF-2_ domain. The figure was generated using PyMol^85^ software.

This study focuses on the PPIs of host G3BP1_NTF2_ protein with the NTD-N_41-174_ of SARS-CoV-2 and emphasizes the inhibition of crucial viral-host protein interactions. The *in-silico* PPIs study predicted the interacting residues of both proteins and revealed the NTD-N_41-174_ derived peptide important for PPIs. After that, the comparative biophysical and biochemical studies supported the participation of residues in molecular interactions of NTD-N_41-174_ and G3BP1_NTF-2_. Primarily, Isothermal Titration Calorimetry (ITC) and fluorescence intensity-based PPIs assays were used to access the binding affinities between purified NTD-N_41-174_ protein, its three different mutants, as well as NTD-N_41-174_ derived peptides with the G3BP1_NTF-2_ protein. Then, the effect of selected small molecules on the disruption of these PPIs was also observed. This systematic approach, coupled with cell-based antiviral assays, shows that compounds significantly impaired virus-host interactions, positioning G3BP1_NTF-2_ as a prime target for developing antivirals against SARS-CoV-2. This pioneering report on small molecules as antivirals targeting host proteins involved in specific PPIs with viral proteins significantly advances the development of effective therapeutics against emerging SARS-CoV-2 variants.

## MATERIAL AND METHODS

### Molecular docking of SARS-CoV-2 N-NTD_41-147_ and G3BP1_NTF-2_ protein

To investigate and analyze PPIs between SARS-CoV-2 N-NTD_41-147_ and G3BP1_NTF-2_ protein as indicated in the reported study of XXX et al., the foremost 3-D structure coordinates of SARS-CoV-2 native N-NTD_41-147_ (6M3M)^80^ and native G3BP1_NTF-2_ (4FCJ)^81^ were extracted from the RCSB Protein Data Bank (PDB)^82^. A docked complex of the N-NTD_41-147_ and G3BP1_NTF-2_ was generated using HADDOCK 2.4^83^ with default parameters. The docked complex was used as the reference for all the computational studies and the notation of the interacting residue involved in the PPIs of N-NTD_41-147_ and G3BP1_NTF-2_. After confirming the interacting residues through a HADDOCK study, the binding affinity of the peptide stretch (N_128-138_) which was present at the interface of NTD_41-147_ and G3BP1_NTF-_2 docked complex with G3BP1_NTF-2_ was determined using AutoDock vina.^84^ The results were analyzed using PyMOL^85^ to identify the key interacting residues in the N_128-138_ peptide responsible for the PPIs.

Subsequently, structural alignment for analysis and comparison of the following G3BP- peptide complex structures: G3BP1-nsP3 complex (PDB: 5FW5^75^), G3BP1 complexed with N_1-25_-IDR region of N-protein (PDB: 7SUO^51^), and the docked structure complex of the newly identified N_128-138_ peptide and G3BP1_NTF-2_ was performed to determine the specific molecular interaction at the interface and similarity in these complexes. Lastly, the multiple sequence alignment (MSA) was performed on peptides that have been reported to interact with G3BP1_NTF-2_, which includes the host protein Caprin1 (PDB: 8TH7)^70^ and proteins of other viruses, including nsP3 of SFV (PDB: 5FW5), the N_1-25_ IDR-1 of the N-protein from SARS-CoV-2 (PDB: 7SUO), and SARS-CoV-2 NTD-N_41-174_ derived peptide (N_128-138_) (PDB: 6M3M), all of which interact with the conserved pocket of the G3BP1_NTF-2_ domain. The analysis was done to identify the key residues of the N_128-138_ peptide (GIIWV) that interact with G3BP1_NTF-2_. Interestingly, sequence analysis indicates that the GIIWV motif in the NTD-N_41-174_ is conserved among coronaviruses (Supporting Information, Figure S1).

### Production of NTD-N_41-174_ protein and G3BP1_NTF-2_ protein

The NTD-N_41-174_ was produced and purified by following a prescribed procedure.^86^ Other than this, expression and purification of the G3BP1_NTF-2_ protein was also performed, as described earlier.^55,87^ The eluted fractions of both target proteins were incubated with TEV protease overnight at 4 °C for the assured cleavage of His-tag. RNA contamination was avoided, and successfully pure proteins were obtained by adding RNAase (Thermo Fisher) at 0.2 mg/ml concentration. His-tag cleaved proteins were further buffer exchanged with 20 mM Tris-HCl, pH 8.0, and 200 mM NaCl. The purity of the cleaved NTD-N_41-147_ and G3BP1 proteins was further validated via 12% sodium dodecyl sulphate-polyacrylamide gel electrophoresis (SDS-PAGE) (Supporting Information, Figure S2). The pure fractions of target proteins were pooled and concentrated using an Amicon ultra centrifugal filter, 10-kDa Molecular weight cut-off (MWCO), and then utilized for further experiments.

### Cloning, expression and Purification of GFP-tagged G3BP1_NTF-2_ (G3BP1_NTF-2_-GFP) protein

For fluorescence-based assay, gene fragment of G3BP1_NTF-2_ (1-139 residues) was PCR amplified using oligonucleotide primers 5’- TACAACGGATCCATGGTGATGGAGAAGCCTAGT-3’ with BamHI & 5’- TGGTGCTCGAGCTAACCAAAGACCTCATCTTGGTAT-3’ (reverse) with XhoI at restriction sites. The restriction enzyme-based digestion of PCR product using BamHI and XhoI enzymes was performed, and then the digested product was ligated into the GFP-tagged pET28c vector using DNA ligase. The CaCl_2_ heat shock method was utilized for the transformation process of competent *E. coli* DH5-alpha cells with the recombinant plasmid. The recombinant plasmid was isolated, and DNA sequencing method was used to confirm the gene insert. Subsequently, the recombinant hexahistidine (6xHis)-tagged protein was transformed into E. coli Rosetta (DE3) cells. The *E. coli* Rosetta (DE3) cells transformants were cultured and incubated overnight in agitation at 180 rpm, 37 °C in 10 ml Luria-Bertani (LB) broth supplemented with 35 µg/ml chloramphenicol and 50 µg/ml kanamycin until the optical density at 600 nm (OD_600_) reached upto 0.8. Then, the induction of recombinant G3BP1-GFP protein by 0.35 mM isopropyl -d-1-thiogalactopyranoside (IPTG) at 16°C for 20 h. The cell disruption was performed in lysis buffer (50 mM Tris pH-7.8, 250 mM NaCl, 5% glycerol) using the French press at 21 kpsi. Cell debris was separated using centrifugation at 12,000 rpm for 1.5 h. The 6xHis-tagged G3BP1-GFP protein was purified by loading the supernatant into a nickel-nitrilotriacetic acid (Ni-NTA) column (BioRad, India) pre-equilibrated with buffer. Protein was eluted with 300mM concentrations of imidazole after 10 mM, 20 mM, 40 mM imidazole washes. Bands corresponding to the molecular weight of 45 kDa were observed on SDS-PAGE (Supporting Information, Figure S3). Fractions containing pure protein bands of the G3BP1-GFP protein were concentrated using an Amicon centrifugal filter 30 kDa MWCO millipore. The G3BP1-GFP protein concentration was determined at OD_280_ by the extinction coefficient method. Likewise, three NTD-N_41-147_ mutants {(V134A), (W133A), (WV133134AA) were Polymerase chain reaction (PCR) amplified (Supporting Information, Figure S4, and Table S1), expressed and purified as described protocol (Supporting Information, Figure S5).

### Thermodynamic-based characterization of PPIs using Isothermal Titration Calorimetry (ITC)

Firstly, the ITC experiments were performed at 25°C^88–90^ using MicroCal ITC_200_ microcalorimeter (Malvern, Northampton, MA),^91^ for the evaluation of the thermodynamic parameters target interacting proteins and peptides involved in PPIs. For the execution of ITC experiments, the NTD-N_41-174_ protein, mutants of NTD-N_41-174_, and G3BP1_NTF-2_ proteins were purified (Supporting Information, Figure S2, and S5) and then extensively dialyzed in the HEPES (20 mM HEPES, pH 8.0 and 20 mM NaCl) buffer. Simultaneously NTD-N derived peptide sequences were chemically synthesized and obtained from Biolinkk Pvt Ltd, India, and stocks was prepared in DMSO. Then, to observe the interactions between target proteins, 10 µM of NTD-N_41-174_ (titration cell) was titrated against 10-15 times higher concentration of G3BP1_NTF-2_ protein (syringe), the same procedure was followed for all three NTD-N derived (N_128-138_) peptides and NTD-N_41-174_ mutants. A detailed description of three peptides and mutants SARS-CoV-2 NTD-N_41-174_ are given below in Table 1.

**Table 1.** Detailed description on peptide sequences of SARS-CoV-2 NTD-N_41-174,_ and Mutants of SARS-CoV-2 NTD-N_41-174_.

| S.No. | Peptides | Sequences |
| --- | --- | --- |
| 1 | N<sub>128-138</sub> peptide 1 | 128KDGIIWVATEG138 |
| 2 | N<sub>128-138</sub> peptide 2 | 128KDGIIAAATEG138 |
| 3 | N<sub>128-138</sub> peptide 3 | 128KAGIIAAATEG138 |

**Mutants of SARS-CoV-2 NTD-N<sub>41-174</sub>**
| S.No. | Mutants | Mutation in residues |
| --- | --- | --- |
| 1 | NTD-N<sub>41-174</sub> mutant 1 | V134A |
| 2 | NTD-N<sub>41-174</sub> mutant 1 | W133A |
| 3 | NTD-N<sub>41-174</sub> mutant 1 | WV133,134AA |

The NTD-N_41-174_ derived peptides (N_128-138_) and titrant inhibitors stock were prepared in DMSO and used after serial dilution in HEPES buffer (20 mM HEPES, pH 8.0 and 20 mM NaCl). Thereafter, to evaluate the inhibition of PPIs by selected compounds, 100 µM of each compound (1:1 concentration ratio) was incubated with 100 µM concentration of G3BP1_NTF-2_ protein for 24 h. Post-incubation, ITC titration reactions were designed by keeping the final titrant concentration (G3BP1_NTF-2_ with incubated compounds in a syringe) set 10 times the analyte (the NTD-N_41-174_ in cell port). Each experiment involved 20 injections (the first injection of 0.5 µL followed by the remaining 19 injections of 2 µL) at time interval of 220 sec with a reference power 10 µcal/sec and a stirring speed of 850 rpm to ensure the homogenous mixing of the solutions. The sample cell and syringe were cleaned after every experiment. The one-site binding model was employed to obtain the binding parameters like reaction stoichiometry (n), enthalpy (ΔH), entropy (ΔS), and association constant (K_A_) using Malvern’s Origin 7.0 Microcal-ITC_200_ analysis software.

### Fluorescence-based biochemical characterization of compounds

The molecular interactions between the NTD-N_41-174_ protein, its above-mentioned mutants, and three N_128-138_ peptides of the SARS-CoV-2 N-protein with the GFP-tagged G3BP1_NTF-2_ protein was determined using a fluorescence-based PPIs assay.^92–94^ The assay was performed using a Synergy HTX multimode plate reader (Agilent BioTek) equipped with Gen5 software. High binding affinity 96-well black fluotrac (CHIMNEY-655076) microplates (Greiner bio-one) were used for the PPIs experiments. Initially, the wells of the microplate were coated with varying concentrations range of the NTD-N_41-174_ protein (0 µM, 10 µM, 25 µM, 50 µM, 100 µM) in duplicate, which were diluted in binding buffer A (0.1 M carbonate-bicarbonate, pH 9.6). It was followed by overnight incubation at 4°C to allow the binding of coated protein to the bottom of wells. The wells were washed twice to remove the unbound protein with washing buffer B (20 mM HEPES, pH 8.5, 20 mM NaCl, 0.05% Tween 20). The NTD-N_41-174_ protein-bound wells were blocked by adding blocking buffer C (20 mM HEPES buffer, pH 8.5, 20 mM NaCl, 1% BSA) and incubated at 25°C for 1 h to stop the non-specific binding. Subsequently, the wells were washed with buffer B to remove the unbounded blocking agents. The purified G3BP1_NTF-2_-GFP protein (Supporting Information, Figure S3) was then added to the wells with bound NTD-N_41-174_ protein to allow the interaction of both proteins, and their fluorescence readings were recorded and analyzed (Supporting Information, Figure S6) after washing wells twice with buffer B to remove the unbound GFP-G3BP1_NTF-2_ protein. In parallel, the purified C-terminal Domain of N protein (CTD-N-purified in our lab) was taken as the negative control because it does not interact with G3BP1_NTF-2_. The plate was incubated for 2 h and rewashed twice with washing buffer B. Then, fluorescence intensity readings were analyzed to get the optimized concentrations of the interacting proteins (Supporting Information, Figure S7). The above-mentioned procedure was again repeated to analyze the interaction of three different N_128-138_ peptides and then three NTD-N_41-174_ mutants with the G3BP1-GFP protein. Further, the pre-incubated 25 µM of G3BP1_NTF-2_-GFP protein with a variable range of selected inhibitors (0 µM, 5 µM, 10 µM, 25 µM, 50 µM) were added to the wells coated with the NTD-N_41-174_ protein to record the fluorescence and analyze PPIs inhibition. The fluorescence intensity was measured using a Synergy HTX multimode plate reader at an excitation wavelength of 485/20 nm and an emission wavelength of 528/20 nm. The data was analyzed using GraphPad Prism^95^ (version 9.0). The statistics signify the average from a duplicate set of reactions with the included standard deviation (SD).

### Cell viability assay

Vero cells were used to propagate the virus and for *in vitro* antiviral studies. Maintenance of the cell was executed using Dulbecco’s modified Eagle’s medium (DMEM; Himedia, India) augmented with 10% fetal bovine serum (FBS; Gibco, USA), 100 units of penicillin, and 100 µg streptomycin/mL (Himedia, India).

MTT (3-(4, 5-dimethyl thiazolyl-2)-2,5-diphenyltetrazolium bromide) assay was employed to determine the viability of Vero cells in the presence of two-fold dilutions of compounds in DMEM supplemented with 2% FBS as described previously.^96,97^ The monolayer of Vero cells in a 96-well plate was incubated with the 2-fold dilution of compounds for 48 h at 37 °C and 5% CO_2_ in a humidified incubator. After incubation, 20 µl of 5mg/ml of MTT was added per well. After 4 h of incubation with MTT, media was discarded, and 150 µl of DMSO was added per well. The absorbance was recorded at 570 nm by Multiskan SkyHigh Microplate Reader (Thermo Fisher), and the percentage viability was calculated by considering the reading of solvent-treated cells as a positive control. The assay was performed in triplicate, and the CC_50_ (50% cytotoxic concentration) was calculated from the dose-response curve in GraphPad Prism 8.

### *In vitro* antiviral assay

As mentioned previously,^98^ of the SARS-CoV-2/Human/IND/CAD1339/2020 strain (GenBank accession no: MZ203529) virus was utilized to perform the antiviral assay. The antiviral activity of compounds was assessed by evaluating the virus inhibition in compound-treated cells compared to virus control (compound non-treated) with the help of quantitative reverse transcription polymerase chain reaction (qRT-PCR) by following the previously mentioned protocols.^98^ Briefly, 24-well with a monolayer of Vero cells was pre-treated with the 2-fold dilution of compounds below their cytotoxic concentration. Then, after washing with PBS, cells were infected with 0.1 multiplicity of infection (MOI) of SARS-CoV-2 for 1.5 h. Post-infection washes with PBS were given, and cells were incubated with respective compound concentrations for 48 h in DMEM supplemented with 2% FBS. Post incubation, the plate was freeze-thawed, RNA was isolated by HiPurA™ Viral RNA Purification Kit for the qRT-PCR, and the cell lysate was also processed for virus titration by TCID_50_ (Median Tissue Culture Infectious Dose) assay as described previously.^98^

## RESULTS

### Molecular Docking for the specificity analysis of NTD-N_41-174_ derived (N_128-138_) peptide involved in PPIs with the G3BP1_NTF-2_ protein

For the identification of the key residues involved in the PPIs of SARS-CoV-2 N-NTD_41-147_ and G3BP1_NTF-2_, an *in silico* molecular docking approach was employed, and the binding energy (HADDOCK score) of both interacting proteins was determined. Initially, the molecular docking generated complex of N-NTD_41-147_ and G3BP1_NTF-2_ with affinity HADDOCK score = −81.3+/-0.5, revealed the residues involved in PPIs of both the target proteins (Figure 1b). After that, the docking of N-NTD derived (N_128-138_) peptide with G3BP1_NTF-2_ revealed (B.E. = −7.8 kcal/mol) and confirmed the specific involvement of N_128-138_ peptide (128KDGIIWVATEG138 residues) in binding with the G3BP1_NTF-2_ protein (Figure 1c-d). These results highlight the identification of novel N_128-138_ peptide and its active contribution in the PPIs.

The comparative structural analysis of G3BP1_NTF-2_ in complex with the NTD of SARS-CoV-2 N-protein (PDB: 7SUO)^51^ and with non-structural protein 3 (nsP3) of alphavirus (PDB: 5FW5)^54^ confirms that the viral protein, NTD and nsP3 bind to the same pocket in the NTF2- like domain of G3BP1_NTF-2_ host protein (Figure 2a). The FGDF motif of Semliki Forest virus (SFV) (nsP3) (orange cartoon) interacts in the groove of G3BP1_NTF-2_ (Figure 2b), and in the case of SARS-CoV-2, the ITFG motif of NTD makes molecular contacts with G3BP1_NTF-2_ (blue cartoon) (Figure 2d). Thus, different families of viruses exploit and target a conserved molecular interaction site within the G3BP1_NTF-2_ to disrupt the normal stress granules formation function of host G3BP1_NTF-2_. Interestingly, this study has identified an additional sequence 128KDGIIWVATEG138 in the NTD-N_41-174_ protein, which binds in a very similar manner to the sequence motifs of other viruses and interacts with the same surface groove of the G3BP1_NTF-2_ protein (Figure 2c). After identifying the N_128-138_ peptide responsible to interact with G3BP1_NTF-2_ specifically. The MSA of different peptides like from host protein Caprin1 (PDB: 8TH7) and different viruses including nsP3 of SFV, APRITFGGPSD motif the SARS-CoV-2 NTD N_1-25_ N-protein, and newly introduced from current study SARS-CoV-2 NTD-N_41-174_ derived peptide (N_128-138_), that interacts within conserved pocket of G3BP1_NTF-2_ domain. The MSA outcomes confirm the participation and importance of the conserved (N_128-138_ peptide) residues for the PPIs with G3BP1_NTF-2_ (Figure 2e). Considering the importance of conserved residues of G3BP1_NTF-2_ for establishing interactions with different viruses, G3BP1_NTF-2_ is referred to as a potential target for antiviral drug discovery. The molecules identified by Mahajan et al. (2022)^55^ that target the conserved pocket in the G3BP1_NTF-2_ were used in this current study to determine the inhibitory effects against the PPIs of the NTD and G3BP1_NTF-2_ proteins.^55^

### Production of NTD of SARS-CoV-2 nucleocapsid protein (NTD-N_41-147_), G3BP1 protein_NTF-2_, mutant proteins of NTD-N_41-147_ and the GFP-tagged G3BP1 (G3BP1-GFP) protein

Purification of both interacting proteins (NTD-N and G3BP1) was performed via Ni-NTA agarose affinity chromatography. The molecular weight of NTD-N (22 kDa), G3BP1 (20 kDa) and the purity of both proteins was observed on 15% SDS-PAGE (Supporting Information, Figure S1). Similarly, the three different NTD-N_41-147_ (W133A mutant, V134A mutant, and WV133134AA) mutants were purified (Supporting Information, Figure S5). For fluorescence-based PPIs assay, the transformation of G3BP1-GFP recombinant plasmid was into *E. coli* BL21 (DE3) cells followed by the IPTG induction at 16°C for 20 h enhanced the overexpression of the protein. The G3BP1-GFP protein (45 kDa) was purified using Ni-NTA agarose affinity chromatography and analyzed by 12% SDS-PAGE (Supporting Information, Figure S2).

### Binding thermodynamics for PPIs of G3BP1_NTF-2_ with NTD-N_41-174_ protein and NTD-N_41-174_ derived peptides (N_128-138_)

ITC isotherm curves help to access the binding affinities of interacting proteins and evaluate the inhibition of PPIs in the presence of inhibitory compounds.^88,90^ Here, the ITC experiments were carried out to determine the biophysical interactions between the target G3BP1_NTF-2_ and the NTD-N_41-174_ protein. Both the proteins have shown a strong interaction, and their calculated dissociation constant (K_D_) was 5.5 µM (Figure 3a; Table 2).

**Figure 3.**
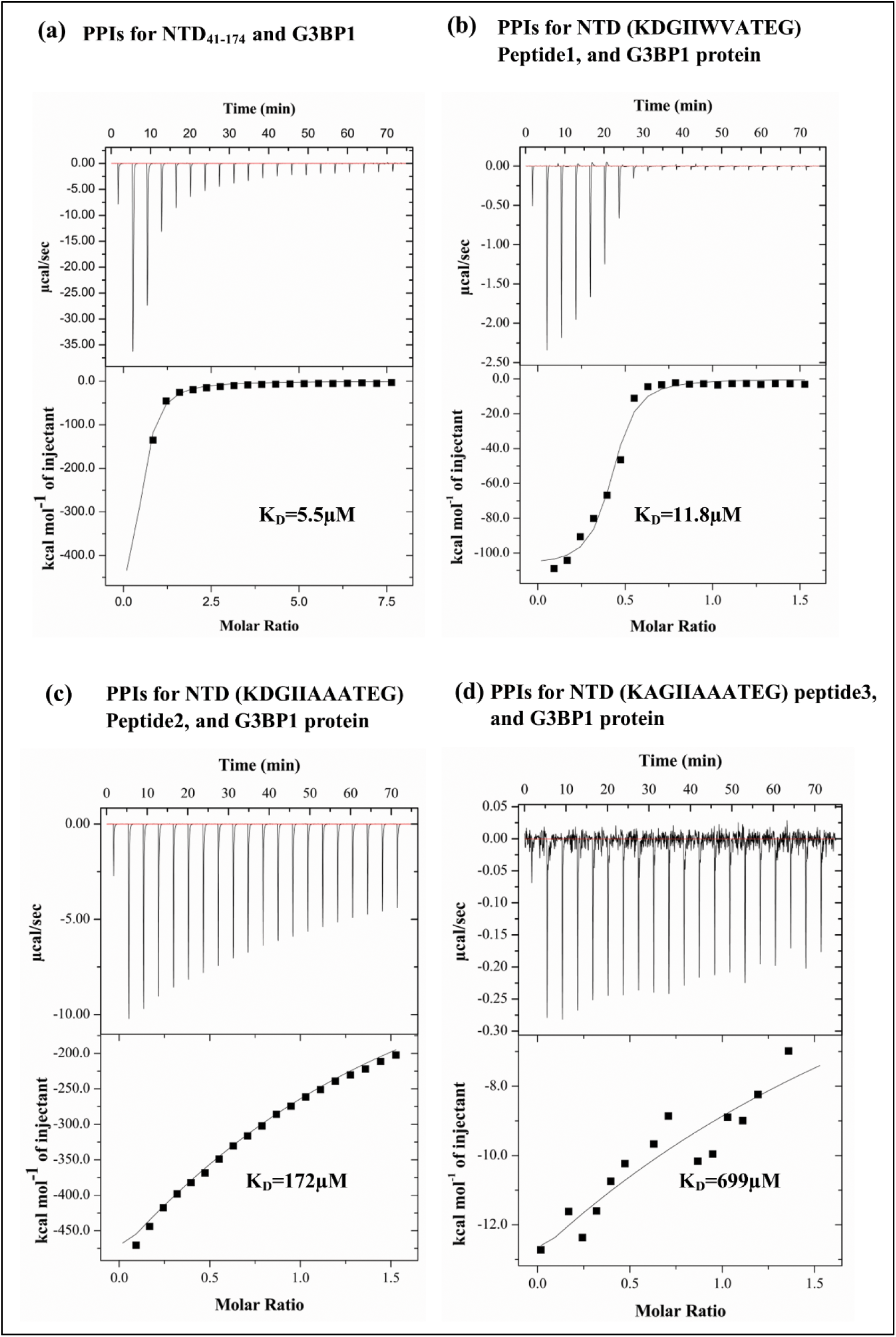
(a) Binding isotherms for PPIs between NTD-N_41-174_ protein and G3BP1_NTF-2_, (b) Binding isotherms for PPIs of N_128-138_ peptide 1 (KDGIIWVATEG), (c) N_128-138_ mutant peptide 2 (KDGIIAAATEG), and (d) N_128-138_ mutant peptide 3 (KAGIIAAATEG) with the G3BP1_NTF-2_ protein using ITC.

**Table 2.** The analysis of thermodynamic parameters for PPIs of three different NTD-N_41-147_ derived (N_128-138_) peptides on titrating against G3BP1_NTF-2_ protein as obtained from ITC are mentioned below.

| Proteins/Peptides<br>for PPIs | n | K<sub>D</sub><br>(μM) | K<sub>A</sub><br>(M<sup>-1</sup>) | ΔH<br>(cal/mol) | ΔS<br>(cal/mol/degree) |
| --- | --- | --- | --- | --- | --- |
| (NTD-N<sub>41-174</sub><br>_G3BP1<sub>NTF-2</sub>)<br>PPIs | 1 | 5.5 | $(1.8)10^5 \pm (1.4)10^4$ | $(-6.9)10^5 \pm (1.3)10^6$ | $(-2.3)10^3$ |
| N<sub>128-138</sub> peptide 1<br>(K**D**GII**W**VATEG)<br>with G3BP1<sub>NTF-2</sub> | 1 | 11.8 | $(8.2)10^4 \pm (6.31)10^3$ | $(-3.3)10^5 \pm (1.2)10^4$ | $(-1.0)10^3$ |
| N<sub>128-138</sub> mutant<br>peptide 2<br>(K**D**GII**A**AATEG)<br>with G3BP1<sub>NTF-2</sub> | 1 | 172 | $(5.7)10^3 \pm (1.3)10^3$ | $(-1.2)10^6 \pm (4.1)10^4$ | $(-4.2)10^3$ |
| N<sub>128-138</sub> mutant<br>peptide 3<br>(K**A**GII**A**AATEG)<br>with G3BP1<sub>NTF-2</sub> | 1 | 699 | $(1.4)10^3 \pm (0.7)10^3$ | $(-9.7)10^3 \pm (1.7)10^3$ | -309 |

Subsequently, the (N_128-138_) peptide 1 (K**D**GII**WV**ATEG), mutant (N_128-138_) peptide 2 (K**D**GII**AA**ATEG), and mutant (N_128-138_) peptide 3 (K**A**GII**AA**ATEG) were titrated against the G3BP1_NTF-2_ protein to analyze the interaction efficiency of the NTD-N_41-174_ derived peptides (N_128-138_). These results demonstrate that the N_128-138_ peptide 1 interacts with G3BP1_NTF-2_ with a dissociation constant (K_D_) of 11.8 µM, showing a relative binding affinity compared to the reference PPIs (NTD-N_41-174_ and G3BP1_NTF-2_, K_D_ = 5.5 µM) (Figure 3a). Titration analysis of NTD-_N41-174_ protein with G3BP1_NTF-2_ in the compared N_128-138_ peptide 1 revealed similar results in PPIs (Figure 3a-b; Table 2). Upon the titration analysis between NTD-N protein and G3BP1 (incubated with NTD-N peptide1), PPIs were observed to be diminished (Supporting Information, Figure S8; Table S2). These findings indicate that the N_128-138_ peptide 1 (KDGIIWVATEG) plays a crucial role in the PPIs between NTD-N_41-174_ and G3BP1_NTF-2_. In contrast, nearly sixteen to sixty folds reduction in PPIs was observed with mutant N_128-138_ peptide 2 and mutant N_128-138_ peptide 3 (containing mutated key interacting residues) (Figure 3c-d) compared to the native N_128-138_ peptide 1 (containing the key interacting residues) (Figure 3; Table 2). The outcomes showed that the residues D129, W133, and V134 are critically involved in the interactions of NTD-N_41-174_ with G3BP1_NTF-2_.

### Binding thermodynamics evaluated the PPIs of NTD-N_41-174_ mutants with G3BP1_NTF-2_ protein

Three different NTD-N_41-174_ mutants were titrated against the G3BP1_NTF- 2_ protein to confirm the participation of key interaction residues involved in PPIs. This study shows that the 128K**DGIIWV**ATEG138 residues of NTD-N_41-174_ play a significant role in the interactions of both the target proteins. The K_D_ values for the double mutants (WV133,134AA) was observed approximately two-four folds higher K_D_ than the single mutants (V134A and W133A) (Figure 4; Table 3). The outcomes of these experiments demonstrated the lower binding affinity of the double mutant as compared to the single residue mutants, which revealed a multi-fold decrease in the PPIs of both target proteins due to double mutation (Figure 4).

**Figure 4.**
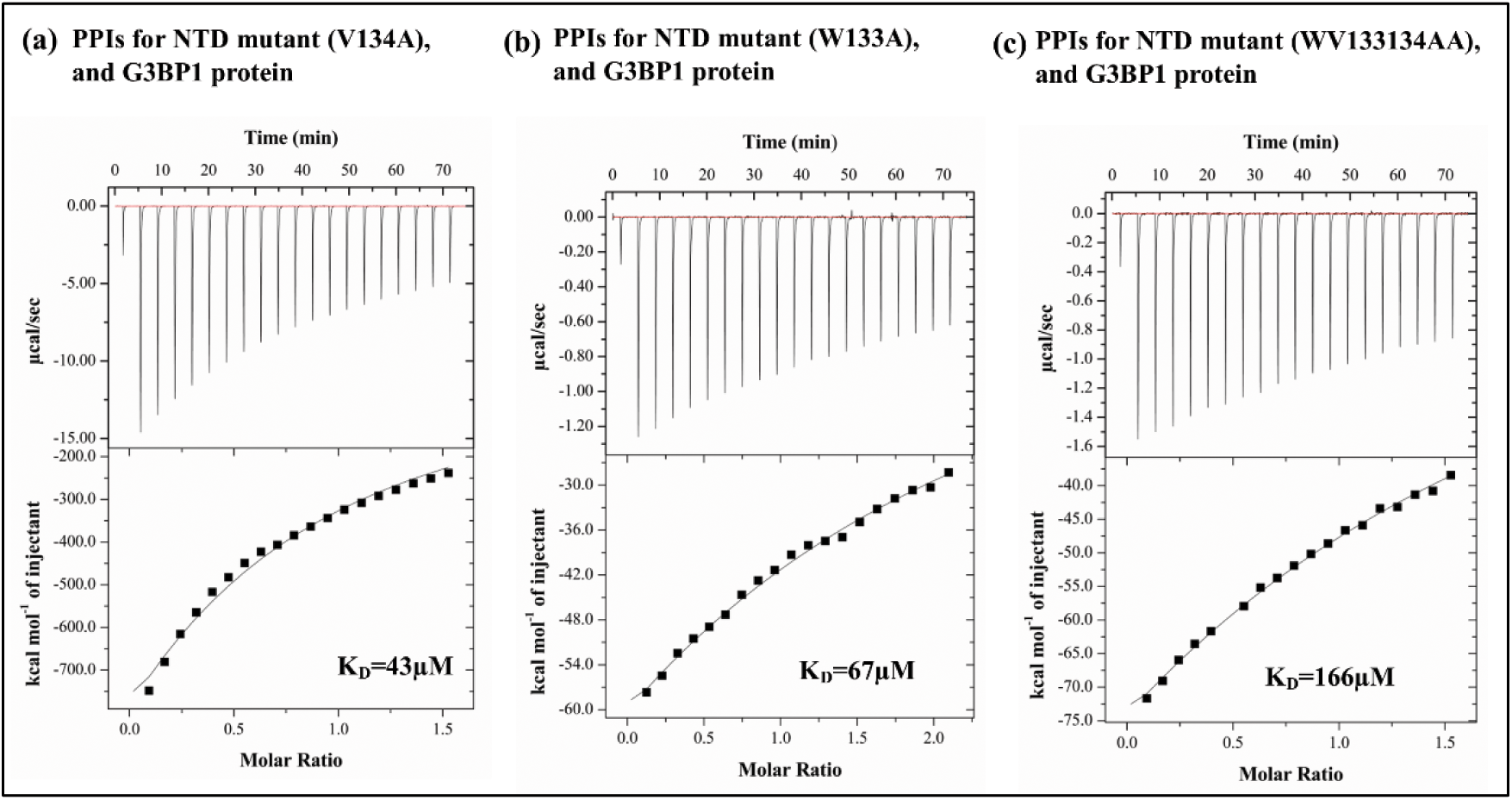
(a) Binding isotherms for PPIs of NTD-N_41-174_ protein mutant (V134A), (b) mutant (W134A), and (c) mutant (WV133134AA) with the G3BP1_NTF-2_ protein measured using the ITC.

**Table 3.** The thermodynamic parameters and analysis for PPIs of NTD-N_41-174_ mutants with G3BP1_NTF-2_ protein was obtained from ITC are mentioned below.

| Mutant proteins for PPIs | n | $K_D$ (μM) | $K_A$ ($M^{-1}$) | $\Delta H$ (cal/mol) | $\Delta S$ (cal/mol/degree) |
| --- | --- | --- | --- | --- | --- |
| NTD-N<sub>41-174</sub> mutant (V134A) with G3BP1<sub>NTF-2</sub> | 1 | 43.6 | $(2.3)10^4 \pm 709$ | $(-3.9)10^5 \pm (0.8)10^4$ | $(-1.2)10^3$ |
| NTD-N<sub>41-174</sub> mutant (W133A) with G3BP1<sub>NTF-2</sub> | 1 | 67.5 | $(1.4)10^4 \pm 640$ | $(-4.5)10^5 \pm (1.5)10^4$ | $(-1.5)10^3$ |
| NTD-N<sub>41-174</sub> mutant (WV133134AA) with G3BP1<sub>NTF-2</sub> | 1 | 166 | $(6.0)10^4 \pm (1.8)10^3$ | $(-4.4)10^6 \pm (1.5)10^5$ | $(-1.4)10^3$ |

### Binding thermodynamics revealed inhibition of NTD-G3BP1_NTF-2_ PPIs

To observe the inhibition of these interacting proteins, NTD-N_41-174_ was titrated against the G3BP1_NTF-2_ protein (pre-incubated with the inhibitors). The selected inhibitors were previously shown to bind with the G3BP1_NTF-2_ protein efficiently^55^. This study shows that these inhibitors efficiently inhibit the interactions between the NTD-N_41-174_ and the G3BP1_NTF-2_ protein. The obtained K_D_ values for the selected inhibitors were several-fold higher than the K_D_ of reference PPIs (Figure 5a), demonstrating adequate inhibition in the interaction of G3BP1_NTF-2_ with NTD-N_41-174_ protein. Interestingly, comparisons of the titration parameters revealed a sharp decrease in the interactions of both target proteins in the presence of inhibitors (Figure 5b-h; Table 4). These results demonstrated the inhibitory capacity of the selected compounds against PPIs of the G3BP1_NTF-2_ and the NTD-N_41-174_ protein.

**Figure 5.**
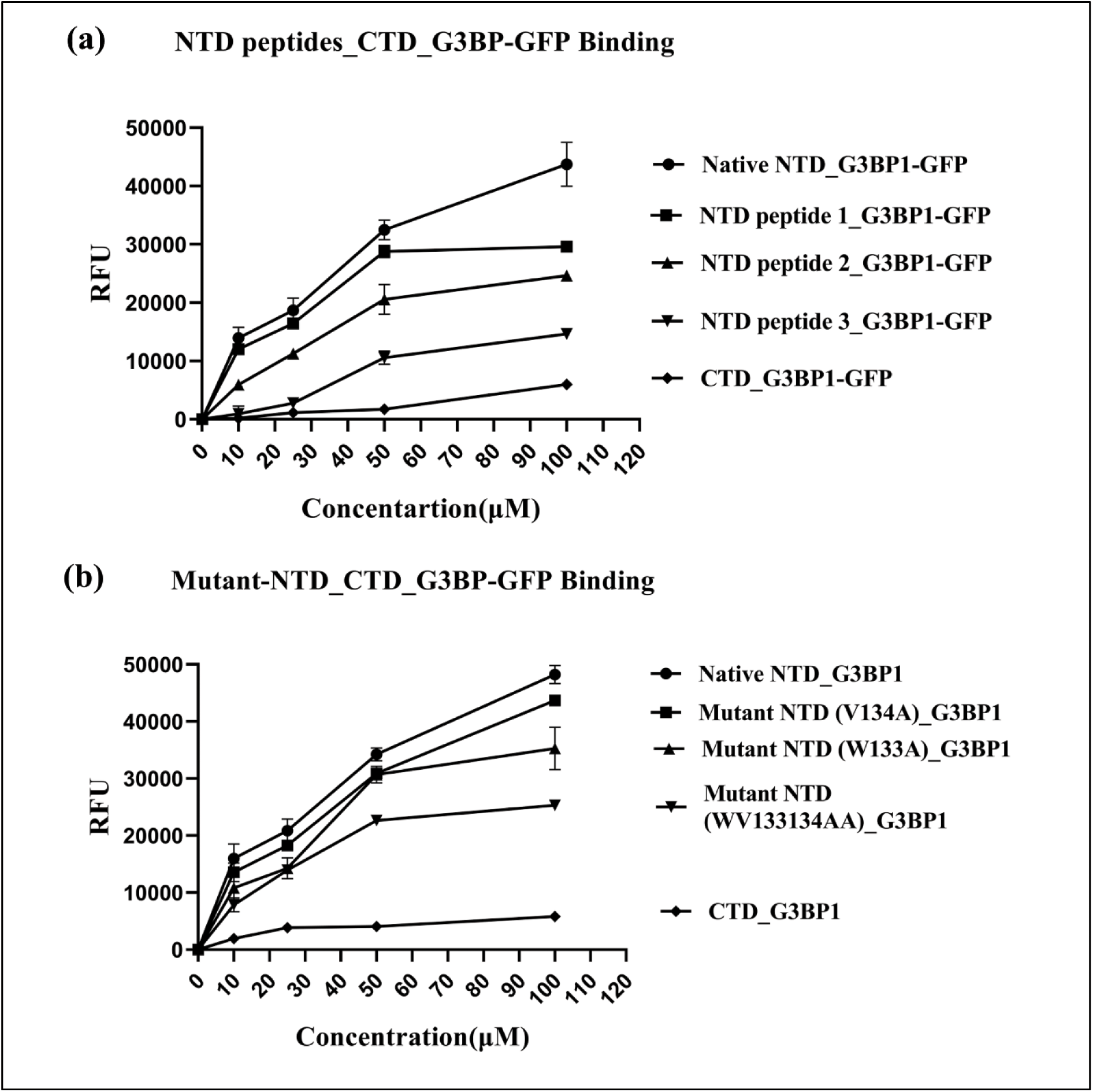
(a) Binding isotherms for PPIs between NTD-N_41-174_ protein and G3BP1_NTF-2_, (b-h) PPIs inhibitions shown by selected compounds using ITC.

**Table 4.** The thermodynamic analysis for PPIs of NTD-N_41-174_ with G3BP1_NTF-2_ was obtained from ITC. The PPIs inhibition parameters in the presence of selected inhibitors are mentioned below.

| Inhibitors | n | K<sub>D</sub><br>(μM) | K<sub>A</sub> (M<sup>-1</sup>) | ΔH (cal/mol) | ΔS<br>(cal/mol/degree) |
| --- | --- | --- | --- | --- | --- |
| (NTD-N<sub>41-174</sub><br>_G3BP1<sub>NTF-2</sub>) PPIs | 1 | 5.5 | (1.8)10<sup>5</sup> ± (1.4)10<sup>4</sup> | (-6.9)10<sup>5</sup> ± (1.3)10<sup>6</sup> | (-2.3)10<sup>3</sup> |
| Fluspirilene | 1 | 234 | (4.2)10<sup>3</sup> ± 491 | (-4.1)10<sup>6</sup> ± (4.3)10<sup>5</sup> | (-1.3)10<sup>4</sup> |
| WIN-62577 | 1 | 76 | (1.3)10<sup>4</sup> ± 841 | (-2.7)10<sup>6</sup> ± (1.3)10<sup>5</sup> | (-9.0)10<sup>3</sup> |
| Naltrindole-hydrochloride | 1 | 58 | (1.8)10<sup>4</sup> ± 986 | (-3.0)10<sup>6</sup> ± (1.3)10<sup>5</sup> | (-1.0)10<sup>4</sup> |
| L-798106 | 1 | 57 | (1.7)10<sup>4</sup> ± (1.0)10<sup>3</sup> | (-4.5)10<sup>6</sup> ± (2.1)10<sup>5</sup> | (-1.5)10<sup>4</sup> |
| GSK1838705A | 1 | 36 | (2.7)10<sup>4</sup> ± (1.6)10<sup>3</sup> | (-2.4)10<sup>6</sup> ± (9.6)10<sup>4</sup> | (-8.0)10<sup>3</sup> |
| SB_222200 | 1 | 25 | (3.8)10<sup>4</sup> ± 682 | (-1.4)10<sup>6</sup> ± (1.6)10<sup>4</sup> | (-4.9)10<sup>3</sup> |
| Dihydroergotamine | 1 | 23 | (4.2)10<sup>4</sup> ± (1.4)10<sup>3</sup> | (-1.6)10<sup>5</sup> ± (1.4)10<sup>4</sup> | -519 |

### Fluorescence intensity-based assessment for PPIs of target NTD-N_41-174_ with GFP- G3BP1_NTF-2_ that validated the inhibitory potential of the selected compounds

Establishing and implementing the fluorescence intensity-based inhibition assay enabled an assessment of the molecular interaction of the NTD-N_41-174_ with the G3BP1_NTF-2_ protein and also determined the inhibitory activity of selected compounds targeting these protein interactions. The measured fluorescence intensity is directly proportional to the PPIs of both the target proteins. For fluorescence-based PPIs assay, PPIs were analyzed by measuring the Relative fluorescence intensity (RFU) for varying concentrations of G3BP1_NTF-2_-GFP (10µM, 25µM, 50µM) against a fix concentration (10 µM) of NTD-N_41-174_ protein (Supporting Information, Figure S6). After optimizing the concentration of the interacting proteins, RFU for the non-binding (a negative control) protein and NTD-N_41-174_ protein at fix concentration was analyzed against the G3BP1_NTF-2_-GFP protein. Finally, the relative binding fluorescence intensity was observed at different concentrations of NTD-N_41-174_ protein (as a binding protein) and CTD-N protein (as a non-binding protein) against the G3BP1_NTF-2_-GFP protein (Supporting Information, Figure S7). After concentration optimization, it was observed that 10 µM NTD-N_41-174_ protein was sufficient to show binding with 25 µM G3BP1_NTF-2_-GFP protein, while no binding was observed between the CTD-N protein and the GFP-tagged G3BP1_NTF-2_ (Supporting Information, Figure S7).

Thereafter, the Fluorescence intensity-based assay validated the involvement of specific residues in PPIs of NTD-N_41-174_ and G3BP1_NTF-2_. The interactions between three different NTD-N_41-174_-derived (N_128-138_) peptides and the G3BP1_NTF-2_-GFP protein produced the results with the progressive decrease in fluorescence intensity in the case of mutant N_128-138_ peptides 2, and mutant N_128-138_ peptides 3 (Supporting Information, Figure 6a). Similarly, the interaction of three different mutants of NTD-N_41-174_ protein with the G3BP1_NTF-2_-GFP demonstrated a same progressive reduction in fluorescence intensity as of NTD-N_41-174_ mutants, particularly in the case of the double mutant (WV133-134AA) NTD-N_41-174_ protein (Supporting Information, Figure 6b). These findings underscore the significance of key interacting residues within the NTD-N_41-174_ protein in mediating its interactions with G3BP1_NTF-2_.

**Figure 6.**
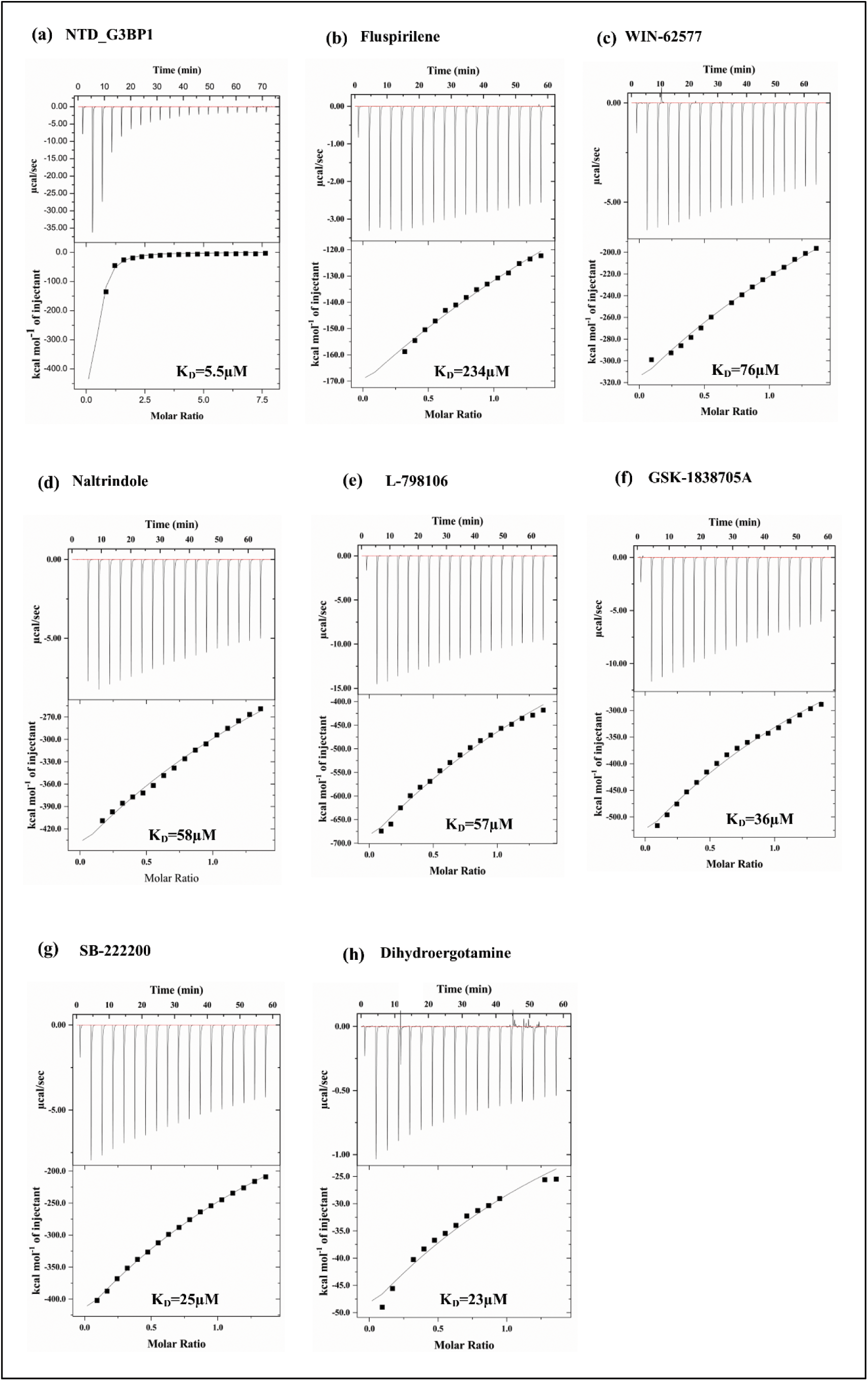
(a) Relative fluorescence intensity v/s concentrations curve for N_128-138_ peptide 1, mutant N_128- 138_ peptide 2, and mutant N_128-138_ peptide 3 with G3BP1_NTF-2_-GFP protein, indicating the optimization of the interaction between N_128-138_ peptide and G3BP1_NTF-2_-GFP protein. N_128-138_ peptide shows strong interactions with G3BP1_NTF-2_ protein and then gradually decreases in interactions on mutating the key interacting residues. (b) Relative fluorescence intensity v/s concentrations curve for NTD-N_41-174_ native (as control) protein, three different mutants of NTD-N_41-174_, and CTD protein (as non-binding protein) with G3BP1_NTF-2_-GFP protein, indicating interactions optimization between NTD-N_41-174_ with the G3BP1_NTF-2_-GFP protein. Three different NTD-N_41-174_ mutants highlight the decrease in PPIs.. The graphs are plotted using GraphPad Prism software. The error bars in the findings show the standard deviation from two measurements.

Later on, to observe the PPIs inhibition, the finalized concentration of 25 µM of G3BP1_NTF-2_- GFP and 10 µM of NTD-N_41-174_ were titrated against different concentrations of the compounds (Supporting Information, Figure S6). Compounds that inhibit PPIs, were shown to reduce the fluorescence intensity by disrupting the formation of protein-protein complexes. This property is essential for drug development and screening of potential drug candidates. The half-maximal inhibitory concentration (IC_50_) of selected seven compounds was calculated using Graph Pad Prism software. The calculated IC_50_ values for different compounds were analyzed and mentioned in Figure 7.

**Figure 7.**
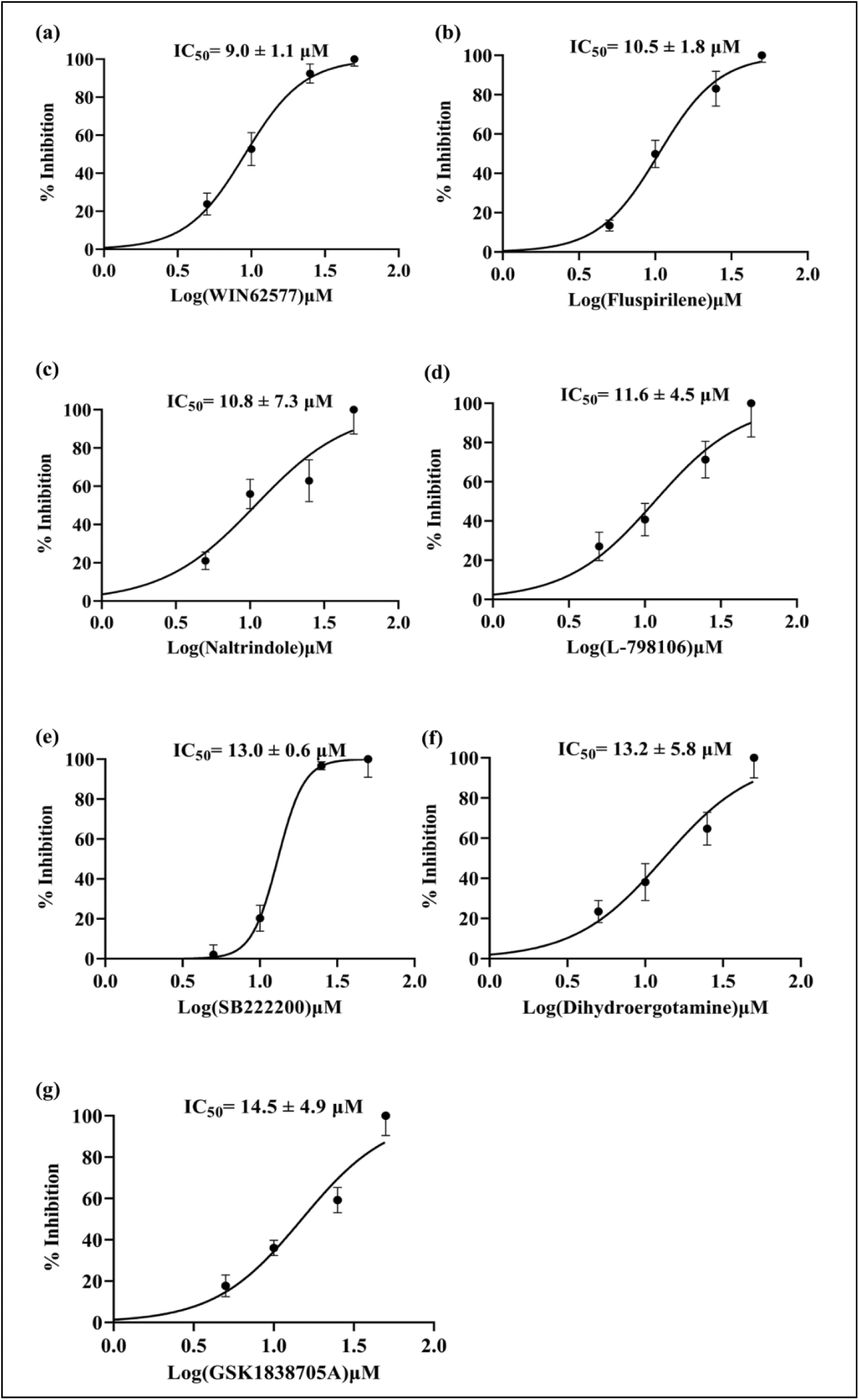
Dose-response curves for the analysis of percentage inhibition for PPIs (NTD-N_41-174_ and G3BP1_NTF-2_) binding versus the concentrations of seven inhibitory compounds plotted in GraphPad Prism software. The error bars in the findings show the standard deviation from two measurements.

### Inhibition of SARS-CoV-2 replication by NTD-G3BP1_NTF-2_ PPI inhibitory molecules

The present study further investigated the capability of selected compounds to deter the replication of SARS-CoV-2 in cell culture. To rule out any effect of compound-induced cellular cytotoxicity, Vero cells were treated with varying micromolar concentrations of compounds for 48 h, and the cellular viability was determined by the MTT assay (Supporting Information, Figure S9). In light of these results, SARS-CoV-2 antiviral assays were performed at non-toxic doses of compounds. Infection inhibition compared to virus control (VC) was quantified in the freeze-thawed cell lysate after 48 h through qRT-PCR and TCID_50_ assay. Among the tested compounds, fluspirilene and WIN_62577 were observed to be the most potent compounds with complete inhibition of SARS-CoV-2 replication at their highest tested concentrations, exhibiting half-maximal effective concentration (EC_50_) values of 1.34 ± 0.03 µM and 1.80 ± 0.0 µM, respectively (Figure 8a; 8c). Both compounds were able to inhibit the replication of SARS-CoV-2 in a dose-dependent manner (Figure 8a; 8c). Similar inhibitory effects were observed when viral titres in harvested cell lysates were enumerated by TCID_50_ assay. At 12.5 µM of fluspirilene, the virus titer was reduced to 10^1^ TCID_50_ /ml compared to the infected non-treated control with 10^5.5^ TCID_50_ /ml (Figure 8b). Interestingly, at a concentration of 3.12 µM of WIN_62577, the virus titer was less than 10^1^ TCID_50_ /ml compared to VC (Figure 8d). The results depict the dose-dependent antiviral behaviour of fluspirilene and WIN_62577 against SARS-CoV-2. The data demonstrates that the compounds fluspirilene and WIN_62577 exhibit potent anti-SARS-CoV-2 activity, providing potential directions for advancing these compounds further along the drug development pipeline.

**Figure 8.**
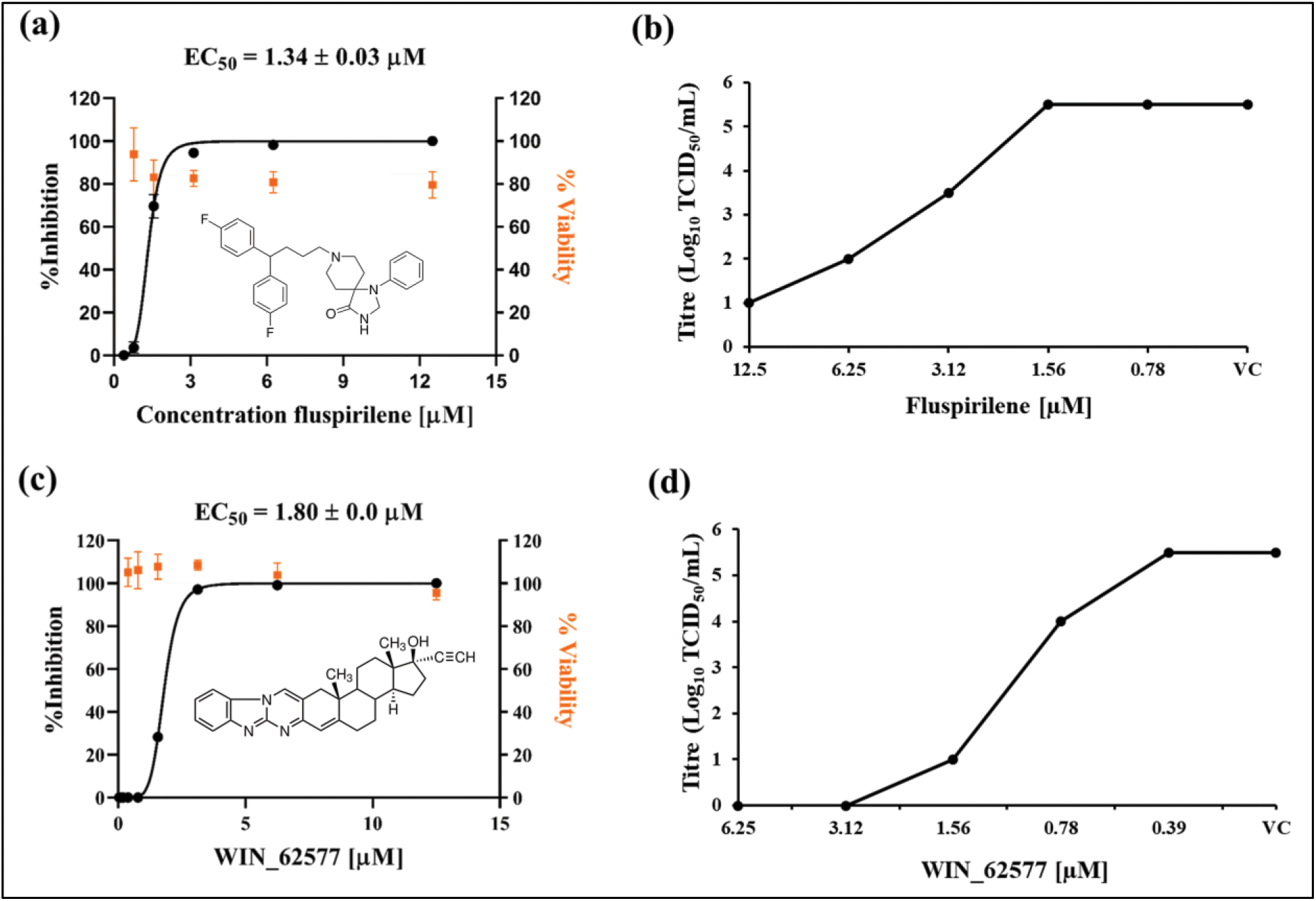
G3BP1_NTF-2_ inhibitors exhibit anti-SARS-CoV-2 activity with minimal toxicity at working concentration. Cellular viability of Vero cells was evaluated in the presence of identified inhibitors by the MTT assay. Data points (orange squares) indicate the mean of 3 replicates in one representative experiment in MTT assay. To assess anti-SARS-CoV-2 activity, infected cells were treated with compounds below their cytotoxic concentration. At the end of the antiviral assay, the viral loads in culture supernatants were quantified by qRT-PCR represented as percentage inhibition relative to the VC (a, and c) and TCID_50_ assay (b, and d). At the highest treated concentration, 4.5-fold and 5.5-fold reduction in virus titre was observed for fluspirilene and WIN_62577, respectively. Cells treated with solvent (DMSO) were used as VC. Sigmoidal dose-response curves (black lines) were fitted, and the EC_50_ values were calculated using GraphPad Prism 8. Data is mean ± SD, n = 2 of biologically independent samples.

## DISCUSSION

The persistent infections of the emergent SARS-CoV-2 strains pose a significant threat to the world’s public health system. It is a matter of concern that more pathogenic, immune evading and highly infectious variants may emerge in the future. Therefore, there is an urgent demand for discovering potent antiviral compounds targeting the virus and host protein interactions critical for viral endurance.^99^ The NTD-N_41-174_ protein of SARS-CoV-2 has a vital role in transcription, genome encapsidation, and virion assembly, as well as its ability to inhibit the host defence mechanism by impeding the G3BP1-mediated SGs formation.^76^ Given the significance of the PPIs in several stages of the viral lifecycle, targeting the interactions between host and viral proteins represents a promising therapeutic strategy beneficial for fighting SARS-CoV-2 and future emerging strains. Therefore, understanding the molecular mechanism underlying the PPIs of NTD-N_41-174_ and G3BP1_NTF-2_ protein is crucial for discovering effective therapeutics against COVID-19 and associated viral infections. Small compounds that can modulate PPIs have proven valuable and advantageous for identifying therapeutics and inhibitors targeting the interactions of NTD-N_41-174_ and G3BP1_NTF-2_ protein, which could form the basis for the host-directed antiviral therapy against SARS-CoV-2. The host protein, G3BP1_NTF-2_, acts as a barrier to viral replication by generating SGs in response to viral infections. However, SARS-CoV-2, among other viruses, has evolved a mechanism to evade host immune responses, as NTD-N_41-174_ protein establishes direct contact with the host’s G3BP1_NTF-2_ protein, promoting viral production by inhibiting the SGs formation. However, the precise molecular basis for the discussed PPIs remains unclear.

The current study introduces a novel NTD-N_41-174_ derived (N_128-138_) peptide that predominantly binds with G3BP1_NTF-2_ and highlights the key interacting residues involved in PPI (Figure 1). The comparative structural analysis of G3BP1_NTF-2_ in complex with the NTD of SARS-CoV-2 N-protein (PDB:7SUO) and with non-structural protein 3 (nsP3) of alphavirus (PDB:5FW5) confirms that the both viral NTD_1-25_ (peptide) and nsP3 protein bind to the same pocket in the NTF2-like domain of G3BP1_NTF-2_ host protein (Figure 2a-b). Thus, different families of viruses exploit and target a conserved molecular interaction site within the G3BP1_NTF-2_ to disrupt the normal stress granules formation function of G3BP1_NTF-2_ (Figure 2c-e). Further, the thermodynamic parameters from ITC experiments identified the binding affinities among the targeted interacting proteins. ITC data revealed the strong molecular interactions between NTD-N_41-174_ and G3BP1_NTF-2_ with a dissociation constant (K_D_) of 5.5 µM (Figure 3a), and inhibition in the presence of selected inhibitors was observed. The biophysical evaluation of PPIs for the three NTD-N_41-174_ derived novel N_128-138_ peptides (Figure 3b-d; Table 2), and three different NTD-N_41-174_ mutants (Figure 4; Table 3) also supported the studies and revealed the importance of the specific residues of N_128-138_ peptide (128KDGIIWVATEG138) for the virus and host PPIs. Thereafter, Inhibition of these PPIs in the presence of seven selected inhibitors was demonstrated (Figure 5b-h; Table 4). All the inhibitors were potentially inhibitory with very high *K_D_* values compared to the *K_D_* of both targeted interacting proteins (Figure 5a; Table 4). A sharp decrease was observed in the binding affinity of G3BP1_NTF-2_ and NTD-N_41-174_ proteins in the presence of inhibitors. These findings indicated the potential of the compounds in inhibiting interactions of the G3BP1_NTF-2_ protein with the SARS-CoV-2 NTD-N_41-174_ protein. To further strengthen the above ITC-based studies, a fluorescence intensity-based protein-protein interactions inhibition assay was further utilized to draw the biochemical assay-based conclusion that revealed the specific concentrations of the proteins were utilized to access the PPIs (Supporting Information, Figure S6; and S7). Simultaneously, the interaction of three different N_128-138_ peptides (Figure 6a) and three different NTD-N_41-174_ mutants (Figure 6b) with G3BP1_NTF-2_-GFP showed the essential residues involved in the PPIs of both target proteins. The fluorescence intensity-based assay showed a gradual decrease in fluorescence intensity, especially with the double mutant NTD-N_41-174_ protein, mutant N_128-138_ peptide 2, and mutant N_128-138_ peptide 3, as compared to the single NTD-N_41-174_ protein mutants and the N_41-174_ derived (N_128-138_) peptide 1, respectively (Figure 6). These results highlight the importance of crucial interacting residues within the NTD-N_41-174_ protein in facilitating its interactions with G3BP1_NTF-2_. After that, the inhibitory activity of the selected compounds against the molecular interactions of the target proteins was determined, and obtained IC_50_ values were in the range of 9 µM to 14.5 µM (Figure 7). The IC_50_ values of selected compounds measured from fluorescence intensity-based assay was a biochemically tested reliable indicator for inhibitory activity against targeted proteins involved in PPIs.

Importantly, the antiviral potential of the compounds that disrupted the PPIs of targeted proteins was assessed by employing *in-vitro* cell culture-based antiviral assessments. WIN-62577 and Fluspirilene were evolved as potential compounds with excellent efficacy in lowering SARS-CoV-2 replication, exhibiting EC_50_ values of 1.8 µM and 1.3 µM, respectively (Figure 8). In an earlier study, Weston et al. (2020) demonstrated that the compound fluspiriline had an IC_50_ of 3.6 µM and a CC_50_ (half-maximal cytotoxic concentration) of 30.33 µM, but they did not report data on its antiviral efficacy, specifically the EC_50_.^100^ Notably, our study has revealed that the two compounds WIN-62577 and fluspiriline exhibited significant antiviral activity with an EC_50_ of approximately 1.8 µM and 1.3 µM, respectively. In conclusion, these findings highlight the importance of targeting the interaction between virus-host proteins and provide valuable insights into developing effective therapeutic strategies that can disrupt PPIs mitigate and alleviate the spread of viral infections.

## ASSOCIATED CONTENT

### Supporting Information

S1: Multiple sequence alignment (MSA) of NTD_41-147_ of N-proteins from four different viruses. S2: Purification of NTD-N_41-174_ protein of SARS CoV-2 and G3BP1_NTF-2_ (Stress granule protein) host protein using Ni-NTA affinity chromatography. Conformation of purified protein on 15% SDS-PAGE gel. S3: 12% SDS-PAGE to confirm the G3BP1_NTF-2_-GFP protein (∼45 kDa) using Ni-NTA affinity chromatography. S4: PCR mutant amplification for NTD-N_41-174_ W133A mutant, V134A mutant, and WV133134AA double mutant. S5: 12% SDS–PAGE analysis of NTD-N_41-174_ W133A mutant, V134A mutant, and WV133134AA double mutant protein purification. S6: Fluorescence-based PPIs assay. Relative fluorescence intensity v/s concentrations curve of NTD protein. S7: Relative fluorescence intensity v/s concentrations curve for NTD (as binding protein) protein and CTD (as non-binding protein) with G3BP1_NTF-2_-GFP protein. S8: Binding isotherms for PPIs of N_128-138_ peptide1 (KDGIIWVATEG) with the G3BP1_NTF-2_ protein using the ITC. S9: The bar graph for the percentage viability of Vero cells after 48 h of compound treatment at indicated different concentrations. Table S1: Primer details of three different NTD-N mutants. Table S2: The analysis of thermodynamic parameters for PPIs of NTD-N peptide with G3BP1 protein as obtained from ITC.

### Accession codes

SARS CoV-2 nucleocapsid protein, UniProtKB P0DTC9 (NCAP_SARS2); and Ras GTPase-activating protein SH3-domain-binding protein 1 (G3BP1), UniProtKB Q13283 (G3BP1_HUMAN).

## AUTHOR INFORMATION

### Authors Contributions

Conceptualization, S.T., P.K., G.K.S., P.K.P. and; methodology, S.T., P.K., G.K.S., P.D., A.S., S.N., and S.C.; Experimentation, P.D., A.S., S.N., and S.C.; formal analysis, P.K., G.K.S., and S.T.; writing-original draft, P.D., A.S., S.N., and S.C., writing-review and editing, P.K., G.K.S., and S.T., supervision, S.T., P.K., G.K.S., and P.K.P.; funding acquisition, P.K., and S.T.; All authors read, revised and approved the manuscript.

## ACKNOWLEDGEMENTS

P.K., G.K.S., and S.T. acknowledge and thank the Science and Engineering Research Board, Department of Science & Technology, Government of India (Project no. IPA/2020/000054) and Scheme for Transformational and Advanced Research in Sciences (STARS)-Ministry of Education (MoE) with (Project ref no: STARS2/2023-0209) for supporting this study. The authors thank the Department of Biosciences and Bioengineering (BSBE) for providing the lab facility, Bioinformatics Centre (BIC), supported by the Government of India (reference number BT/PR40141/BTIS/137/16/2021), the Macromolecular Crystallographic Facility (MCU) for the computer facility. The authors also thank the instrumentation facilities provided by the Institute Instrumentation Centre (IIC) and Ashok Soota Molecular Medicine Facility at the Indian Institute of Technology Roorkee (IIT Roorkee). The authors thank the BSL3 facility at the Indian Veterinary Research Institute, Izatnagar, Bareilly, Uttar Pradesh, India, and the Virus containment facility (BSL-3) at IIT Roorkee for the antiviral experiments on SARS-CoV-2.

## ABBREVIATIONS

PPIs: Protein-protein interactions
G3BP1: Ras GTPase-activating protein-binding protein 1
G3BP1_NTF-2_: NTF-2 like domain of Ras GTPase-activating protein-binding protein 1
NTD: N-terminal domain
SARS-CoV-2: Severe acute respiratory syndrome coronavirus 2
N: Nucleocapsid protein
NTD-N: NTD of SARS-CoV-2 nucleocapsid protein
NTD-N_41-147_: NTD (41-147 residues) of SARS-CoV-2 nucleocapsid protein
N_128-138_: NTD-N_41-174_ derived (N_128-138_) peptide
SGs: Stress granules
EC_50_: effective concentration
RNP: Ribonucleoprotein complexes
IDRs: Intrinsically disordered regions
CTD-N: C-terminal domain
CTD-N, LOPAC^1280^: C-terminal domain of nucleocapsid library of pharmacologically active compounds
CHIKV: Chikungunya virus
nsP3: non-structural protein 3
ITC: Isothermal Titration Calorimetry
SDS-PAGE: Sodium Dodecyl Sulphate-Polyacrylamide gel electrophoresis
G3BP1-GFP: GFP-tagged G3BP1
6xHis: hexahistidine
LB: Luria-Bertani
OD_600_: optical density at 600 nm
Ni-NTA: nickel-nitrilotriacetic acid
n: stoichiometry
ΔH: enthalpy
ΔS: entropy
K_A_: binding association constant
SD: standard deviation
DMEM: Dulbecco’s modified Eagle’s medium
FBS: fetal bovine serum
qRT-PCR: quantitative reverse transcription polymerase chain reaction
MTT: 3-(4, 5-dimethyl thiazolyl-2)-2,5-diphenyltetrazolium bromide
MOI: multiplicity of infection
K_D_: dissociation constant
VC: virus control
RFU: Relative fluorescence intensity.

## ETHICS DECLARATIONS

The authors declare no competing interests.

