## Supplementary material for "Disruption of molecular interactions between G3BP1 stress granule host protein and nucleocapsid (NTD-N) protein impedes SARS-CoV-2 virus replication": Supplimentry file

### ***Supporting information***

#### *Material and methods.*

#### Multiple sequence alignment (MSA) of NTD of N- proteins

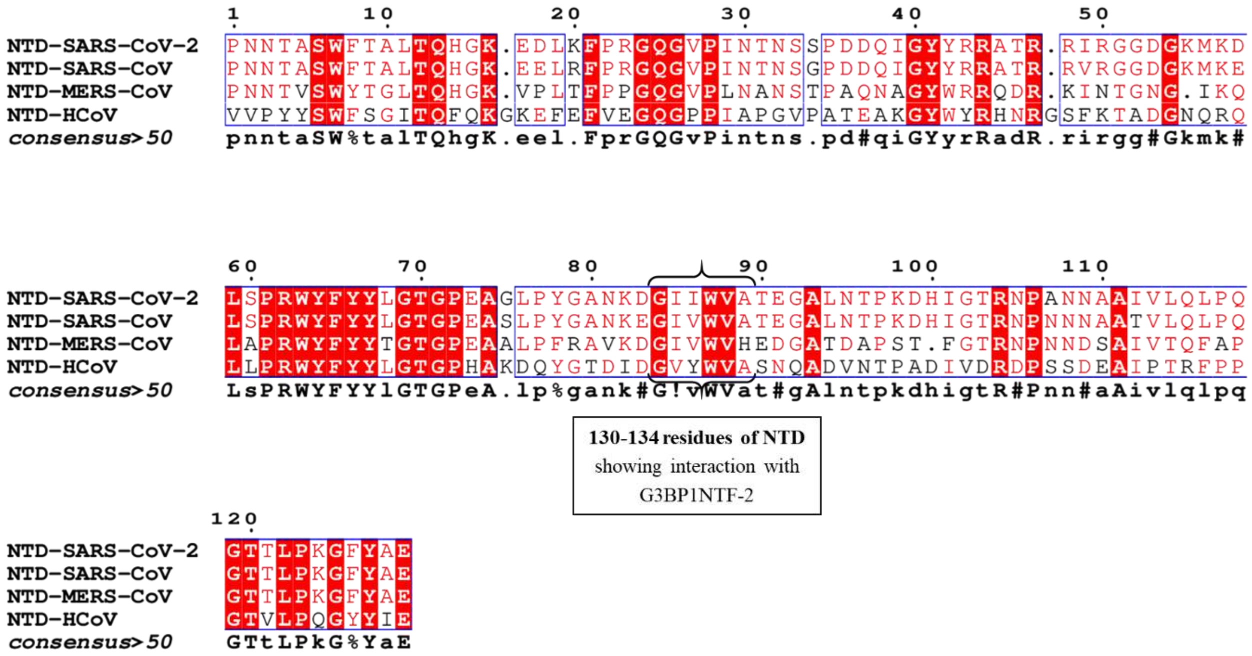


**Figure S1.** Multiple sequence alignment (MSA) of $NTD_{41-147}$ of N- proteins from four different SARS-CoV-2 (NTD-SARS-CoV-2), SARS-CoV (NTD-SARS-CoV), Middle East respiratory syndrome coronavirus (MERS-CoV) NTD-MERS-CoV, and Human coronavirus (NTD-HCoV), these results showed conservation of GIIVW in coronaviruses.

**Purification of NTD$_{41\text{-}174}$ of SARS-CoV-2 nucleocapsid protein (NTD-N), G3BP1$_{NTF\text{-}2}$ protein**

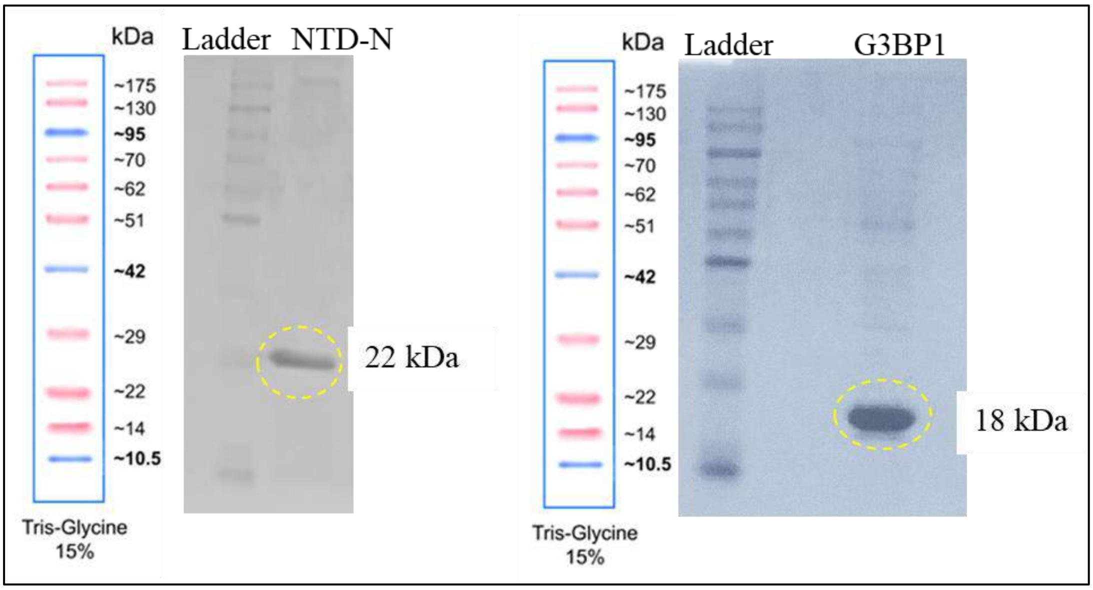


**Figure S2.** Purification of NTD-N protein of SARS CoV-2 and G3BP1 (Stress granule protein) host protein using Ni-NTA affinity chromatography. Conformation of purified protein on 15% SDS-PAGE gel.

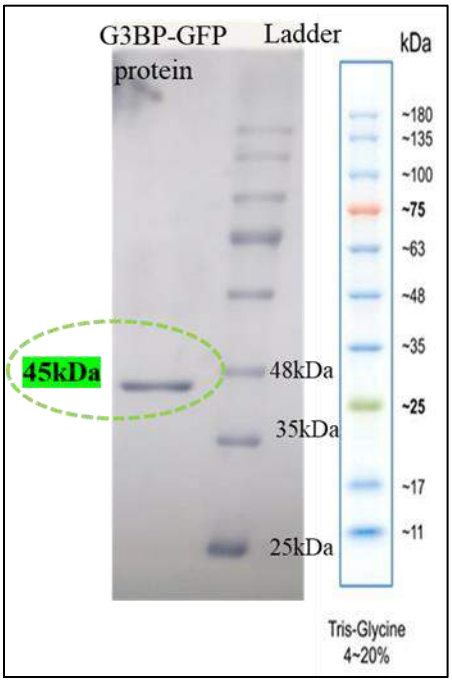

**Figure S3.** 12% SDS-PAGE to confirm the G3BP1-GFP protein (~45 kDa) using Ni-NTA affinity chromatography.

**PCR mutagenesis of NTD-N W133A mutant, V134A mutant, and WV133134AA mutant**

The genomic DNA of NTD-N was used as a template for the NTD-N W133A mutant, V134A mutant, and WV133134AA mutant gene amplification; the oligonucleotide primers used are listed below in table S1 as:

**Table S1.** Primer details of three different NTD-N mutants.

| S. No. | NTD-N mutants | Primers details |
|---|---|---|
| 1. | NTD-N W133A | Forward: 5' CAAAGACGGCATCATAGCGGTTGCAACTGAGGG 3' |
| | | Reverse: 5' CCCTCAGTTGCAACCGCTATGATGCCGTCTTTG 3' |
| 2. | NTD-N V134A | Forward: 5' CGGCATCATATGGGCTGCAACTGAGGGAGC 3' |
| | | Reverse: 5' GCTCCCTCAGTTGCAGCCCATATGATGCCG 3' |
| 3. | NTD-N WV133134AA | Forward:5' CTAACAAAGACGGCATCATAGCGGCTGCAACTGAGGGAGCCTTG 3' |
| | | Reverse:5' CAAGGCTCCCTCAGTTGCAGCCGCTATGATGCCGTCTTTGTTAG 3' |

The mutant gene was amplified and digested DPN I into a pET-28c vector and then transformed into E. coli DH5α cells and the positive clone harbouring the recombinant plasmid were further confirmed by restriction digestion and DNA sequencing.

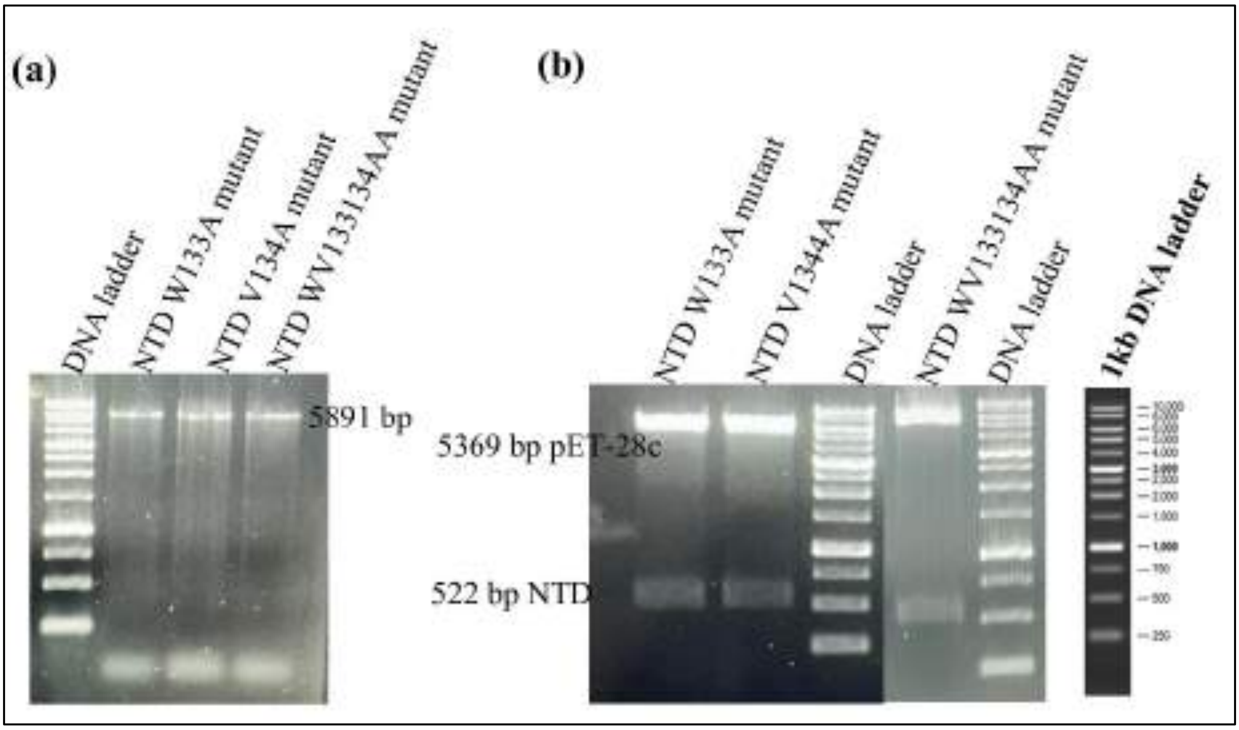


**Figure S4.** (a) PCR mutant amplification for NTD-N W133A, V134A, and WV133134AA. (b) Restriction enzymes digestion of expression vectors for W133A, V134A, and WV133134AA mutant's plasmid digested with Nde I + Xho I.

**Purification of SARS-CoV-2 NTD-N W133A mutant, V134A mutant, and WV133134AA mutant**. The confirmed mutants was then transformed into E. coli BL21 (DE3) cells for protein expression. The gene expression was induced with 0.2 mM isopropyl β-d-thiogalactopyranoside (IPTG) at 16 °C for 18 h. Mutant protein purification of NTD-N was performed via Ni-NTA agarose affinity chromatography. The elution buffer containing 250 mM imidazole and buffer (50 mM Tris-HCl, 500 mM NaCl, and pH 7.0) was used to elute the protein. The molecular weight of NTD-N (21.7 kDa), and mutant proteins size and purity was observed on 12% SDS-PAGE (Figure S4).

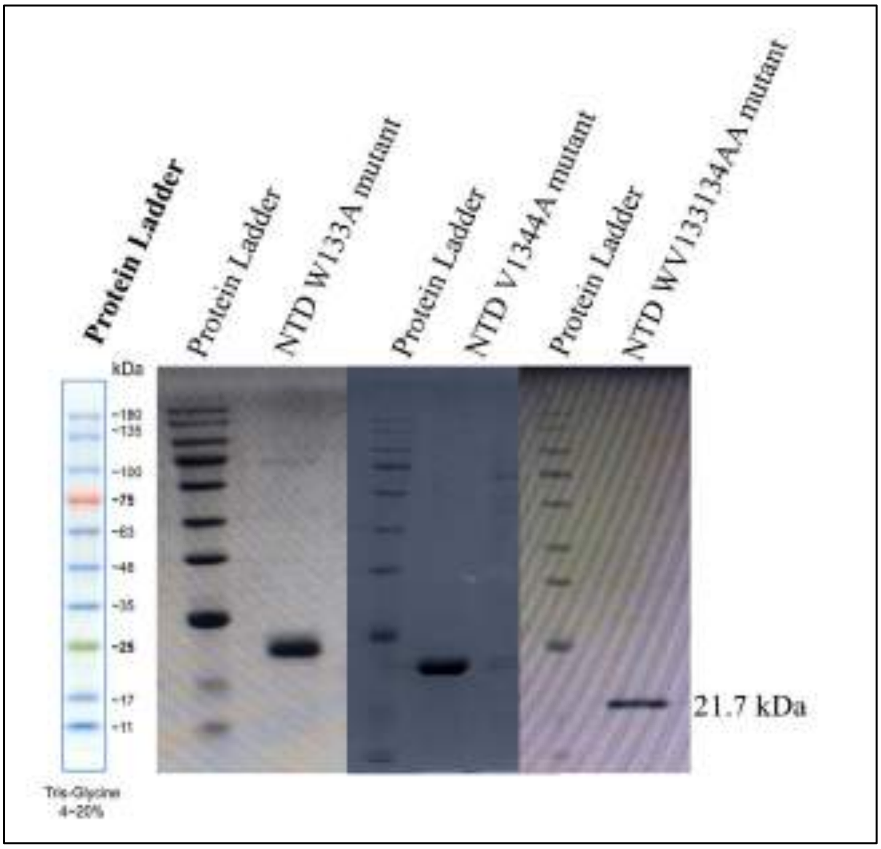


**Figure S5.** 12% SDS–PAGE analysis of NTD-N W133A, V134A, and WV133134AA mutant protein purification with labelled protein marker and represented the purified mutant protein bands of 21.7 kDa.

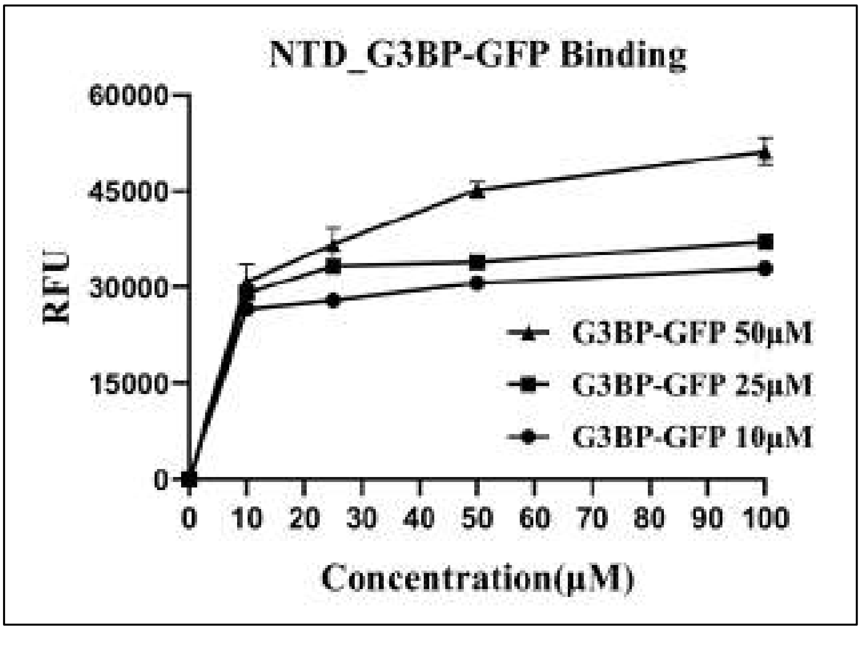


**Figure S6.** Fluorescence-based PPIs assay. Relative fluorescence intensity v/s concentrations curve of NTD protein, indicating interactions optimization of both proteins at different concentrations of G3BP1-GFP (0μM, 10μM, 25μM, 50μM), and keeping NTD concentration constant (10 μM).

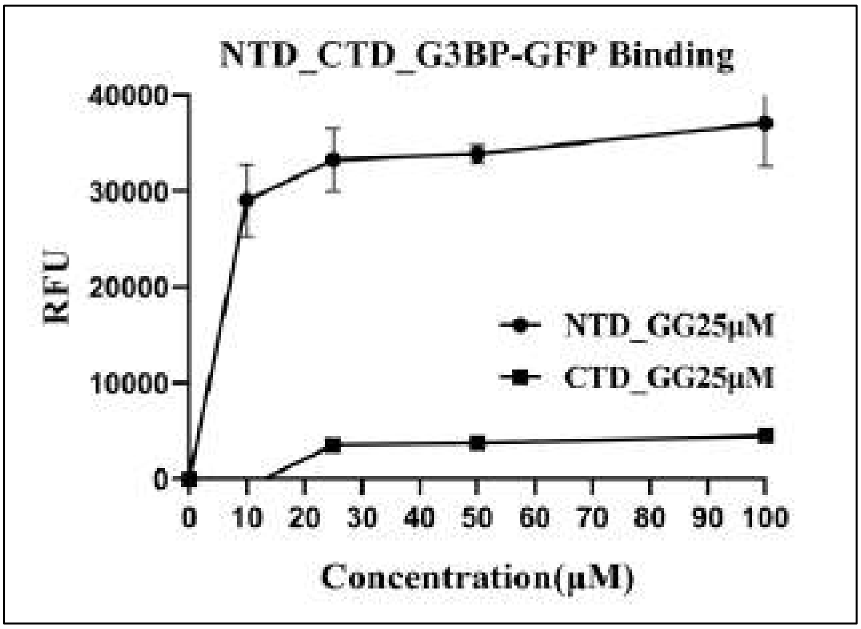


**Figure S7.** Relative fluorescence intensity v/s concentrations curve for NTD (as binding protein) protein and CTD (as non-binding protein) with G3BP1-GFP protein, indicating interactions optimization of NTD (10 μM) with G3BP1-GFP (25 μM).

**Binding thermodynamics analysis of PPIs of NTD-N peptide with G3BP1 protein:** The NTD-N peptide (KDGIIWVATEG) was titrated against the G3BP1 protein to analyse the interaction efficiency of the NTD-N peptide. This study shows that the NTD-N peptide binds with the G3BP1 protein ($K_D$ = 12 μM) with the relative efficiency to the reference PPIs (NTD-N and G3BP1, $K_D$ = 5.5 μM) (Figure ; Table ). Upon the titration analysis between NTD-N protein with G3BP1 (incubated with NTD-N peptide1), PPIs was observed to be diminished (Figure S8 b; Table ST2). Interestingly, these observations revealed that the NTD-N peptide (KDGIIWVATEG) participates in the NTD-N and G3BP1 proteins.

**Table S2.** The analysis of thermodynamic parameters for PPIs of NTD-N peptide with G3BP1 protein as obtained from ITC are mentioned below.

| Proteins for PPIs | n | $K_D$ (μM) | $K_A$ ($M^{-1}$) | ΔH (cal/mol) | ΔS (cal/mol/degree) |
|---|---|---|---|---|---|
| NTD-N peptide (KDGIIWVATEG) with G3BP1 | 1 | 12 | $(8.2)10^4$ ± $(6.31)10^3$ | $(-3.3)10^5$ ± $(1.2)10^4$ | $(-1.0)10^3$ |

| | | | | | |
|---|---|---|---|---|---|
| NTD-N protein with G3BP1 (incubated with NTD-N peptide1) | 1.4 | 755 | $(1.3)10^3 \pm (4.2)10^2$ | $(-4.9)10^4 \pm$ $(1.2)10^4$ | -146 |

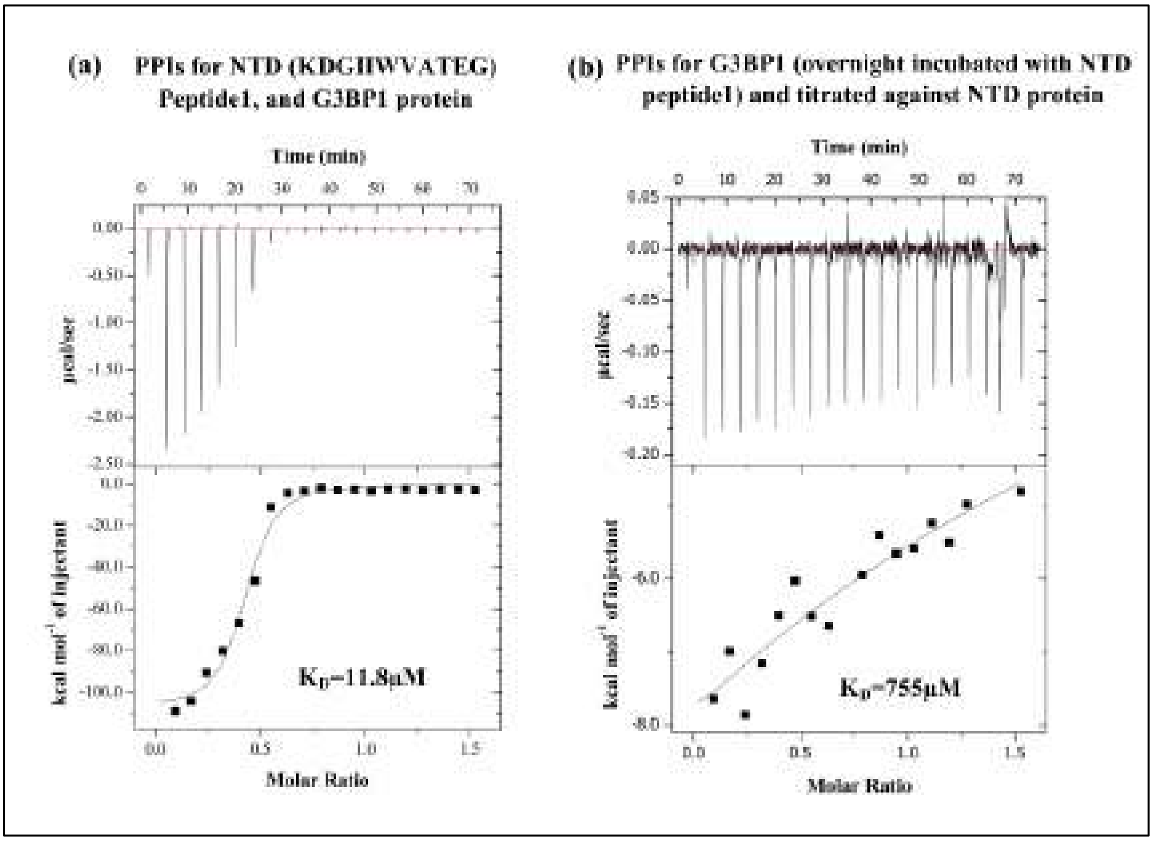


**Figure S8.** (a) Binding isotherms for PPIs of NTD-N peptide1 (KDGIIWVATEG) with the G3BP1 protein, and (b) PPIs inhibition measured after incubating the NTD peptide1 overnight with G3BP1 protein and titrated against NTD protein using the ITC.

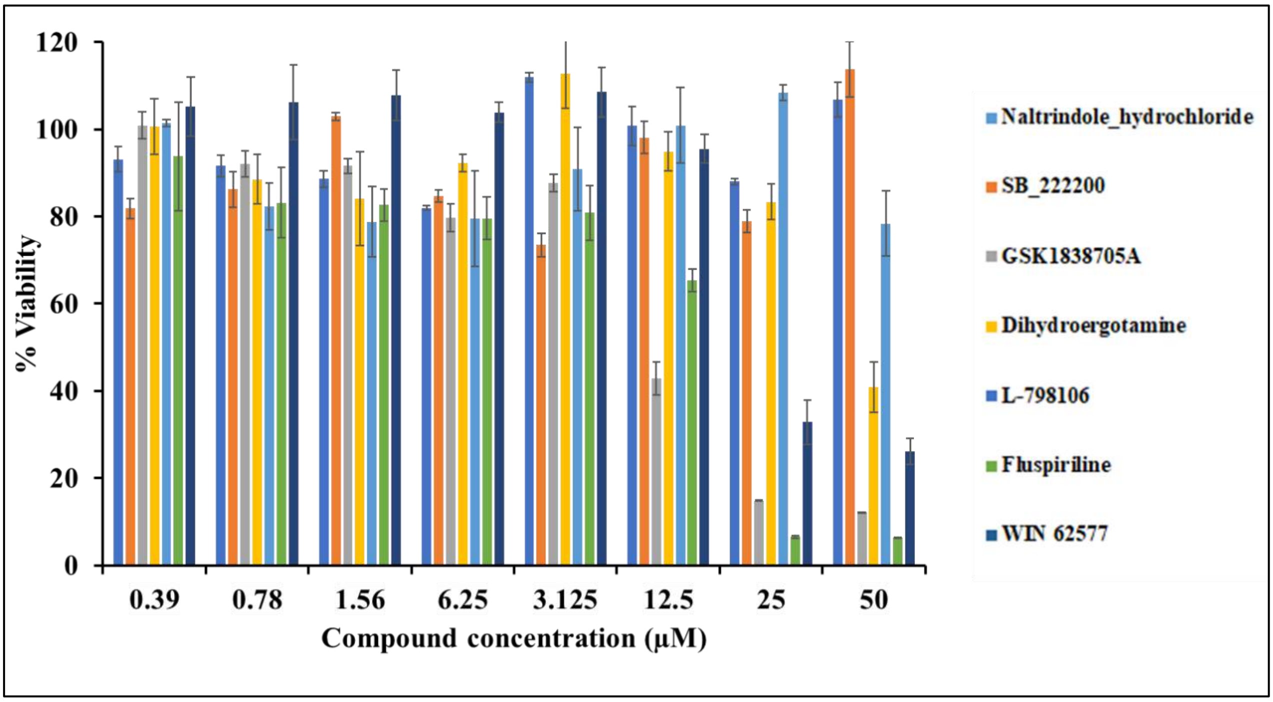


**Figure S9.** The bar graph represents the percentage viability of cells after 48 h of compound treatment at indicated different concentrations. X and Y-axes correspond to the micro molar range compound concentrations and percentage viability, respectively. The viability was calculated by considering the solvent-treated cells as a control. The data was analyzed using Microsoft Excel.
